# RB1 loss triggers dependence on ESRRG in retinoblastoma

**DOI:** 10.1101/2021.12.15.472842

**Authors:** Matthew G. Field, Jeffim N. Kuznetsoff, Michelle G. Zhang, James J. Dollar, Michael A. Durante, Yoseph Sayegh, Christina L. Decatur, Stefan Kurtenbach, Daniel Pelaez, J. William Harbour

**Affiliations:** Bascom Palmer Eye Institute, Sylvester Comprehensive Cancer Center, and Interdisciplinary Stem Cell Institute, University of Miami Miller School of Medicine, Miami, FL 33136, USA

## Abstract

Retinoblastoma (Rb) is a deadly childhood eye cancer that is classically initiated by inactivation of the RB1 tumor suppressor. Clinical management continues to rely on nonspecific chemotherapeutic agents that are associated with treatment resistance and toxicity. Here, we analyzed 103 whole exomes, 16 whole transcriptomes, 5 single-cell transcriptomes, and 4 whole genomes from primary Rb tumors to identify novel Rb dependencies. Several recurrent genomic aberrations implicate estrogen-related receptor gamma (ESRRG) in Rb pathogenesis. RB1 directly interacts with and inhibits ESRRG, and RB1 loss uncouples ESRRG from negative regulation. ESRRG regulates genes involved in retinogenesis and oxygen metabolism in Rb cells. ESRRG is preferentially expressed in hypoxic Rb cells *in vivo*. Depletion or inhibition of ESRRG causes marked Rb cell death which is exacerbated in hypoxia. These findings reveal a novel dependency of Rb cells on ESRRG, and they implicate ESRRG as a potential therapeutic vulnerability in Rb.

## Introduction

Retinoblastoma (Rb) is the most common pediatric eye cancer and an important cause of childhood cancer death worldwide (*1*). Despite considerable improvements in treatment and patient survival over the past century (*2*), severe visual impairment and loss of the eye are still common, due in part to treatment resistance and toxicity associated with currently used chemotherapeutic agents. A better understanding of molecular dependencies in Rb could lead to more specific and effective targeted therapies. Rb is almost always initiated by mutational inactivation of the *RB1* tumor suppressor in a susceptible retinal progenitor cell (*3, 4*), and it is generally thought that subsequent aberrations are required for full malignant transformation (*5, 6*). Several recurrent genomic events have been described in Rb, including mutations in *BCOR* and *CREBBP,* and chromosome copy number variations (CNVs) affecting 1q, 2p, 6p, 13q and 16q (*7–12*). However, there has been limited functional validation and no clinically relevant targeted therapies associated with any proposed secondary drivers. Here, we searched for novel drivers of Rb progression by performing a large integrative multi-omics analysis of data from whole-exome sequencing (WES), whole-genome sequencing (WGS), RNA-sequencing (RNA-seq), single-cell RNA sequencing (scRNA-seq), and other methods. We discovered that the estrogen-related receptor gamma (ESRRG) is an essential mediator of hypoxic adaptation and cell survival in Rb that is constitutively activated by RB1 loss and is subsequently affected by recurrent genomic aberrations in Rb. These findings suggest a selective pressure to increase ESRRG dosage during Rb progression and offer a potential target for therapy.

## Results

### Genomic landscape of retinoblastoma implicates ESRRG in tumor progression

WES data from 103 primary Rb samples were analyzed (**Fig. 1** and **fig. S1**). Mutations were detected in *RB1* in 94% of samples, including a stopgain in 41 cases (39.8%), isodisomy of chromosome 13q in 36 cases (35.0%), loss of heterozygosity of *RB1* in 19 cases (18.5%), splicing mutations in 20 cases (19.4%), homozygous deletion of *RB1* in 9 cases (8.8%), and chromothripsis involving the *RB1* locus in 9 cases (8.8%). Other recurrent mutations were found in *BCOR* (17%), *FCGBP* (5%), *NSD1* (5%), *BRCA2, CREBBP, DST, MACF1*, and *PI4KA* (4% for each), *SCN5A* (3%), and *CHD1, NCOA3*, and *PML* (2% for each). The most common CNVs included 1q gain (67%), 6p gain (66%), 16q loss (56%), 2p gain (44%), 16p loss (29%), and 7q gain (26%). The *MYCN* locus at 2p24 was amplified in 6 cases, 3 of which had no detectable *RB1* mutation. We next searched for functional patterns in the genomic landscape using Ingenuity Pathway Analysis (*13*). Strikingly, most of the proteins encoded by recurrently mutated genes participate in an estrogen receptor/estrogen-related receptor (ESR/ESRR) interaction network involved in development, neurogenesis, metabolism, hypoxia, and cell cycle regulation (**Fig. 2A**). *ESRRG* was the only ESR/ESRR family member that was highly expressed by RNA-seq among 16 primary Rb tumor samples (**Fig. 2B**). Likewise, *ESRRG* was the predominant ESR/ESRR family member expressed by scRNA-seq in 5 primary Rb tumors (**Fig. 2C-E and fig. S2**). *ESRRG* is located on chromosomal region 1q41, which undergoes arm-level gain in two-thirds of Rb cases; 2 of 4 samples analyzed by WGS revealed local chromosomal rearrangements upstream of or involving the *ESRRG* locus (**fig. S3**). These findings suggest that mutations and genomic aberrations that affect *ESRRG* are common in Rb.

**Fig. 1.**
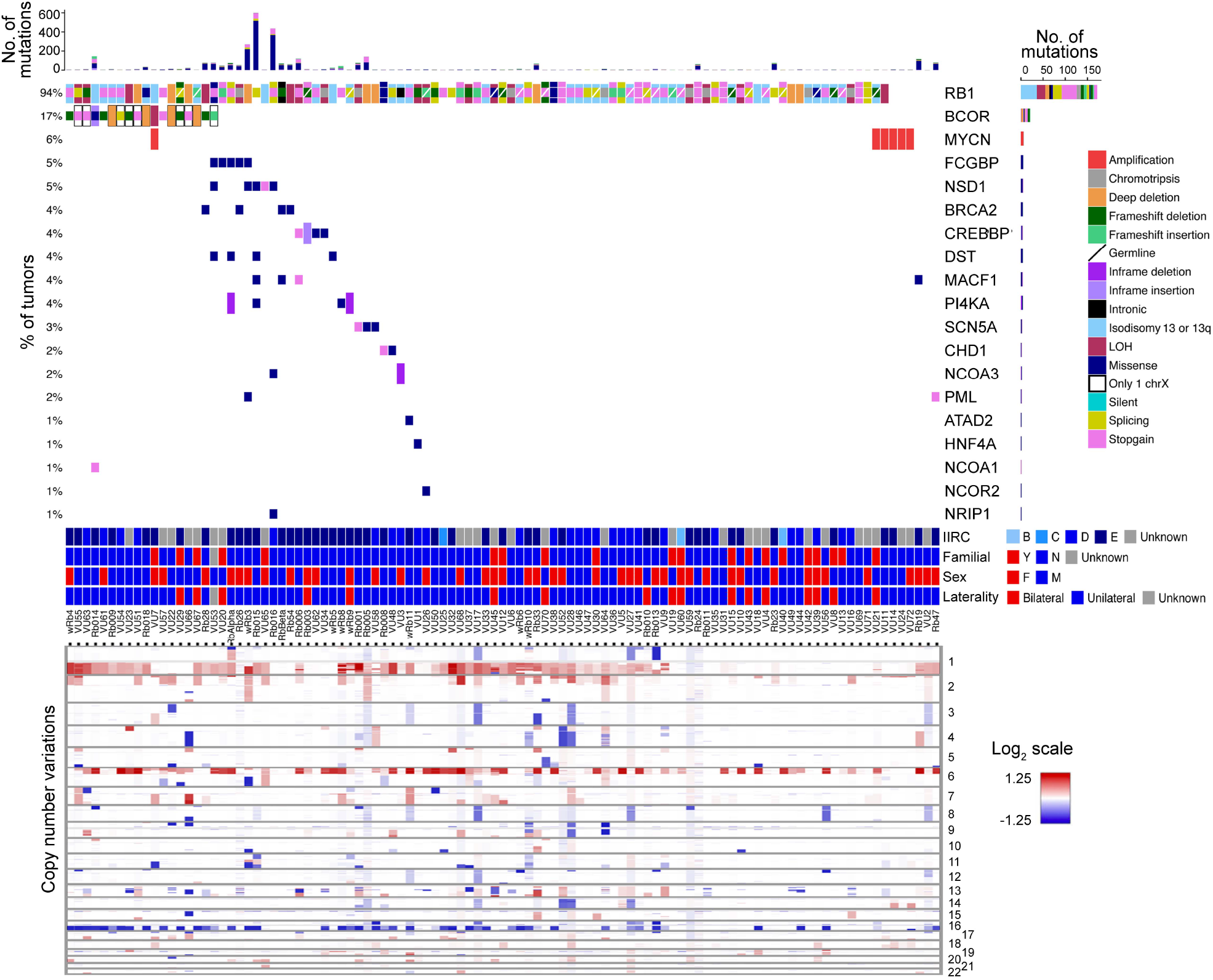
Genomic landscape of 103 primary retinoblastomas. Oncoprint of 103 primary retinoblastoma samples analyzed by whole exome sequencing. Data include status of the most commonly mutated genes, types of mutations, International Intraocular Retinoblastoma Classification (IIRC) status, family history of retinoblastoma, gender, laterality, and chromosome copy number aberrations.

**Fig. 2.**
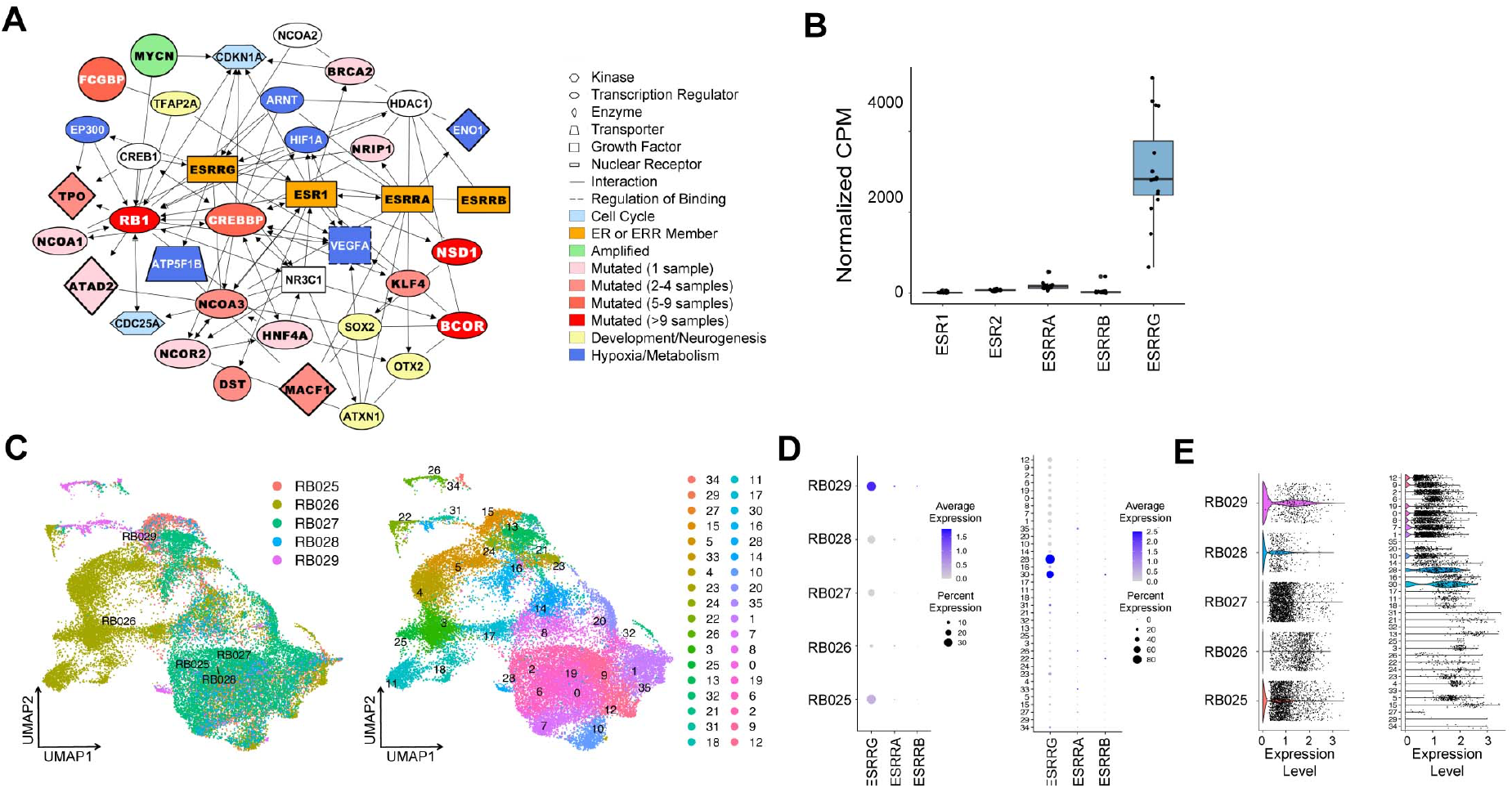
Recurrent genomic aberrations point to ESRRG as a common target for deregulation in retinoblastoma. (**A**) Ingenuity pathway analysis of the most common mutations in 103 retinoblastoma samples. (**B**) RNA-seq analysis of mRNA expression for ESR/ESRR family members in 16 primary retinoblastoma tumors. (**C**) UMAP plots of scRNA-seq data from 5 primary retinoblastoma tumor samples obtained at enucleation, sorted by tumor sample (left) and Seurat cluster (right). (**D**) Dot plot analysis of scRNA-seq data showing relative single-cell expression of ESRRA, ESRRB and ESRRG by tumor sample (left) and Seurat cluster (right). (**E**) Violin plots of scRNA-seq data showing ESRRG expression by tumor sample (left) and Seurat cluster (right).

### ESRRG regulates genes involved in retinogenesis and oxygen metabolism in Rb cells

To gain further insight into how aberrant ESRRG regulation may drive Rb progression, we performed chromatin immunoprecipitation using an anti-ESRRG antibody followed by DNA sequencing (ChIP-seq) in our RB006 and RB018 low-passage Rb cell lines (**fig. S4A**). A high-confidence dataset of 8163 significantly enriched peaks shared between both RB006 and RB018 were used for downstream analyses (**fig. S4B**). ESRRG was enriched at ERRE transcriptional regulatory motifs, as expected, but also at binding motifs associated with retinal and neuronal transcription factors CRX, OTX2, NEUROD, and LHX family members (**fig. S4C**). Interestingly, many ESRRG peaks occurred in regulatory regions containing one or more of these other binding motifs but not an ERRE motif, suggesting that ESRRG can interact with chromatin independently of its canonical binding site. Using peak location analysis, ~15% of ESRRG peaks occurred in promoter regions (± 3kb from the TSS), whereas almost half occurred at presumed enhancer regions between 10kb and 100kb from the TSS, with ESRRG peaks occurring at 446 (24.3%) of 1834 annotated retinal/neural super-enhancer regions (**fig. S4C**)(*14*).

To correlate ESRRG chromatin localization with gene expression, we performed RNA-seq before and after depletion of ESRRG in RB006 and RB018 cells, and in two more of our recently established Rb cell lines (RB016 and wRB6). Of the 8,163 significant ESRRG ChIP-seq peaks, 1,226 were associated with 738 differentially expressed genes (FDR<0.05), including 249 up-regulated genes (353 peaks) and 489 down-regulated genes (873 peaks). Upregulated genes were enriched for pathways related to cell cycle regulation and oxidative phosphorylation, whereas down-regulated genes were enriched for pathways related to neurogenesis, hypoxia, and estrogen signaling. CRX, OTX2, NEUROD, and LHX binding motifs were enriched in both up- and down-regulated genes, with 18.0% of peaks located in promoter regions (± 3kb from the TSS) and 52% located 10-100kb from the TSS (**Fig. 3A-D** and **fig. S5**). Consistent with these findings, scRNA-seq analysis demonstrated a significant correlation between Rb cells expressing *ESRRG* and those expressing *CRX* and *OTX2* (**fig. S6**).

**Fig. 3.**
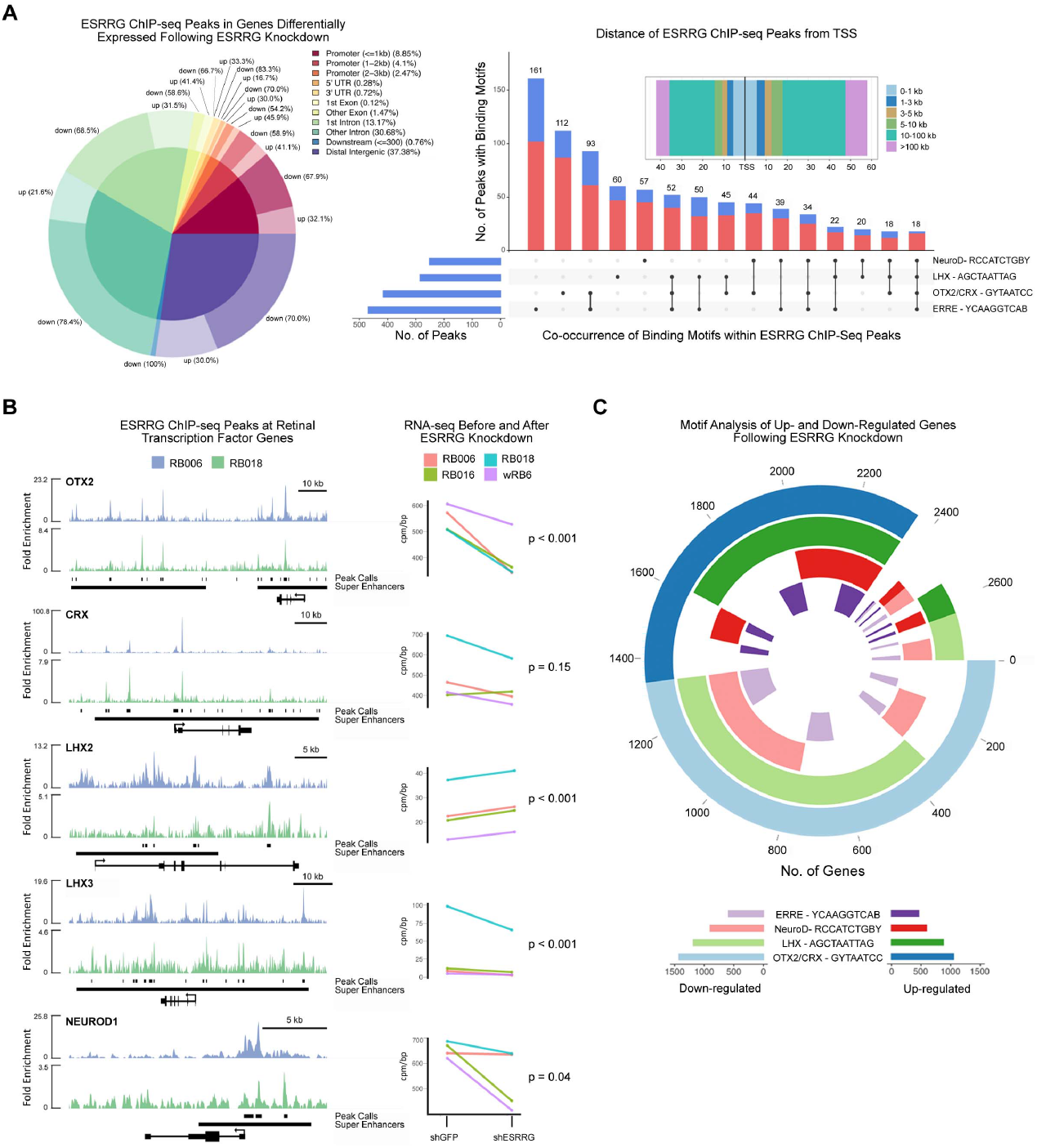
ESRRG regulates genes involved in retinogenesis and oxygen metabolism in retinoblastoma. (**A**) Integrated analysis of ESRRG ChIP-seq peaks and RNA-seq differentially expressed genes with or without shRNA-mediated knockdown of ESRRG in recently established retinoblastoma cell lines. ChIP-seq was performed in RB006 and RB018 cells, and RNA-seq in RB006, RB016, RB018, and wRB6 cells. The pie chart displays the percentage of peaks located within gene regions that are associated with differentially expressed genes (inner circle) or up-regulated and down-regulated genes (outer donut). The rectangular plot (top right) exhibits the distance of the peaks from the transcription start site (TSS). The bar plot shows presence and cooccurrence of the most significantly enriched transcription factor binding motifs (FDR < 0.05) found within the ESRRG ChIP-seq peaks associated with up-regulated (blue) or down-regulated (red) genes after ESRRG knockdown. (**B**) ESRRG ChIP-seq track plots in RB006 cells (blue) and RB018 cells (green) at key retinal transcription factors involved in retinal development and differentiation. Peak calls from the pooled datasets, super enhancer locations, exon and intron locations, and direction of transcription are indicated below peak plots. Corresponding RNA-seq data for each are shown for RB006 (red), RB016 (green), RB018 (blue), and wRB6 (purple) retinoblastoma cells engineered to express shRNA directed against ESRRG (shESRRG) or control (shGFP). P-values were calculated after batch and dispersion correction using EdgeR. (**C**) Circos plot showing presence and co-occurrence of ERRE (purple), NeuroD (red), LHX (green), and CRX/OTX2 (blue) binding motifs within ±3kb from the TSS of all significantly up-regulated (darker colors) and down-regulated (lighter colors) genes (FDR < 0.05) following ESRRG knockdown in RB006, RB016, RB018, and wRB6 cells (pooled data shown). Individual genes are depicted as radii around the circle, and each donut represents the indicated binding motifs that are present ±3kb from the TSS of each gene. The numbers on the outside of the circle increase counterclockwise and represent the number of significantly altered genes recognized by Homer motif analysis.

### RB1 interacts with and inhibits ESRRG

One of the frequently mutated genes implicated in the ESR/ESRR interaction network was *RB1* itself (**Fig. 2A**). ESR1, ESRRA, and ESRRG contain a VXXLYD motif, similar to the IXXLFD RB1-binding motif in E2F1 and E2F2 (**Fig. 4A**)(*15–18*). We found that RB1 and ESRRG coimmunoprecipitate and that this interaction is disrupted by alteration of the VXXLYD motif in ESRRG (**Fig. 4B**). Among the 3 ESR/ESRR family members with a VXXLYD motif, only ESRRG contains this motif within its DNA binding domain (**Fig. 4A**), raising the possibility that RB1 inhibits ESRRG’s transcriptional activation by blocking its interaction with ERRE regulatory motifs (*19*). Consistent with this idea, ectopic expression of wildtype ESRRG (V5-ESRRG-WT) activated a luciferase reporter downstream of the ERRE promoter, and this activation was enhanced by shRNA-mediated knockdown of RB1 in RB1-wildtype HEK293 cells (**Fig. 4C**). Expression of an ESRRG mutant in which the VXXLYD motif was mutated (V5-ESRRG-MUT) resulted in greater activation of the reporter, which was only minimally enhanced by RB1 knockdown, indicating that loss of the RB1-ESRRG interaction results in constitutive ESRRG activity. Further supporting this finding, the ERRE luciferase reporter was activated by wildtype ESRRG in RB1-null C33A cells, and this activation was abrogated by ectopic expression of RB1. The RB1 binding mutant V5-ESRRG-MUT also activated the reporter, and it was resistant to inhibition by RB1. Consistent with previous work in prostate cancer cells (*20*), we found that ectopic expression of V5-ESRRG-WT in HEK293 cells induced p21 expression associated with hypophosphorylation (activation) of RB1, whereas knockdown of endogenous ESRRG resulted in depletion of p21 and hyperphosphorylation (inactivation) of RB1 (**Fig. 4D**). ESRRG was enriched at the *CDKN1A* gene (encoding p21), and knockdown of ESRRG decreased *CDKN1A* RNA expression in Rb cell lines, indicating direct transcriptional activation of *CDKN1A* by ESRRG (**Fig. 4E**). Thus, RB1 directly inhibits ESRRG, which in turn negatively regulates itself by activating p21. After *RB1*, the second most commonly mutated gene in Rb is the co-repressor *BCOR* (**Fig. 1**), and we found that BCOR represses ESRRG transcriptional activity independently of RB1 (**fig. S7**), suggesting that mutations in this gene also serve to increase ESRRG activity.

**Fig. 4.**
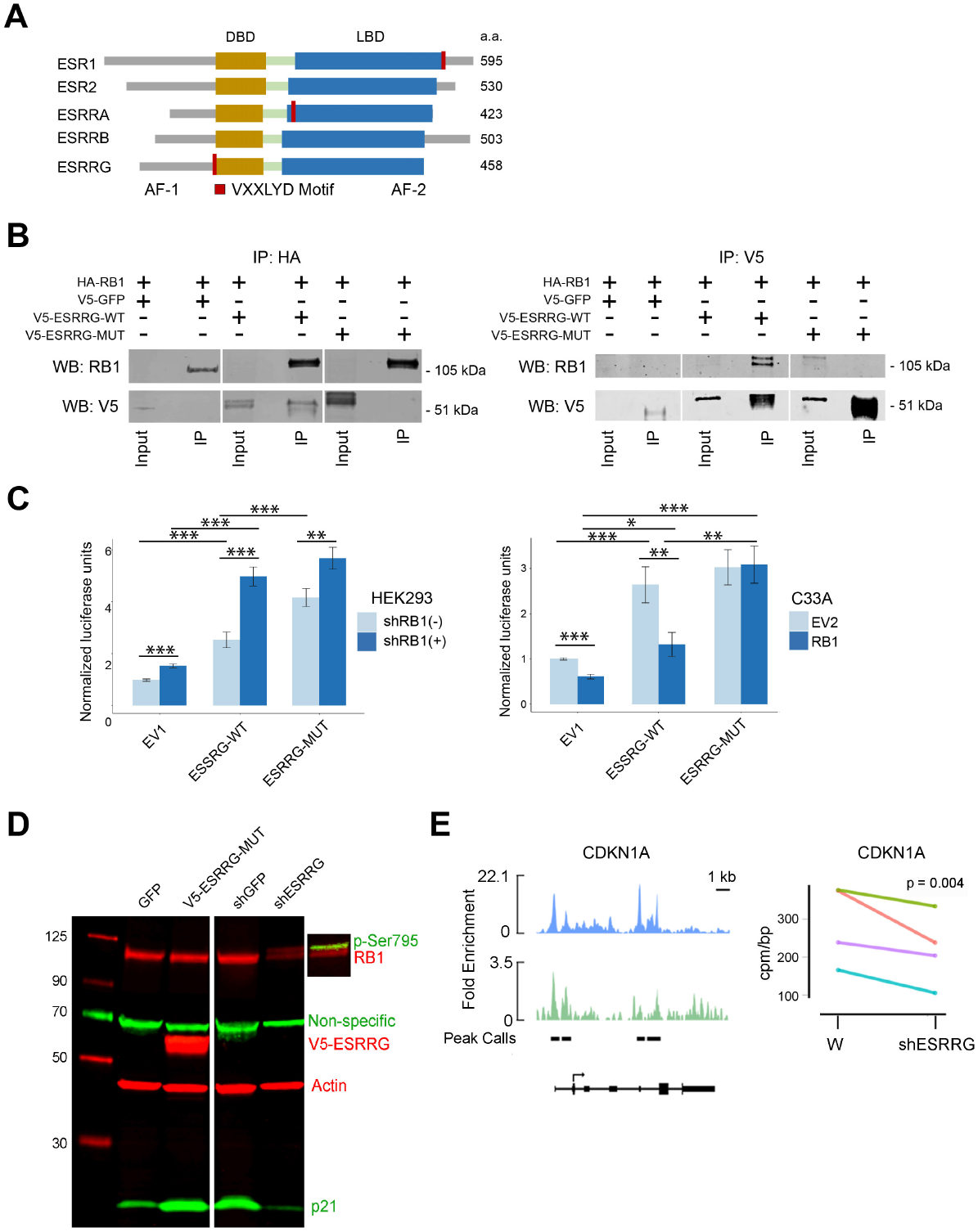
RB1 interacts with and inhibits ESRRG. (**A**) Protein maps showing location of RB1-binding VXXLYD motifs in estrogen receptors (ESR1 and ESR2) and estrogen-related receptors (ESRRA, ESRRB, and ESRRG). AF-1 and AF-2, activation function domains 1 and 2, respectively; a.a., amino acids. (**B**) Western blot (WB) for ectopically expressed HA-tagged RB1 (HA-RB1), V5-tagged wildtype ESRRG (V5-ESRRG-WT), or ESRRG with mutated VXXLYD motif (V5-ESRRG-MUT) in RB1-null C33A cells following immunoprecipitation for the HA tag (left). Western blot for ectopically expressed HA-RB1, V5-ESRRG-WT, or V5-ESRRG-MUT in RB1-null C33A cells following immunoprecipitation for the V5 tag (right). (**C**) Normalized ERRE luciferase reporter activity with or without ectopic expression of V5-ESRRG-WT or V5-ESRRG-MUT, and with or without shRNA-mediated depletion of RB1 (shRB1+ and shRB1-, respectively) in HEK293 cells (top). Normalized ERRE luciferase reporter activity with or without ectopic expression of V5-ESRRG-WT, V5-ESRRG-MUT, and RB1 in C33A cells (bottom). EV1 and EV2, empty vector controls. *, P<0.05; **P<0.01; ***, P<0.001. (**D**) Western blot in HEK293 cells following ectopic expression of V5-ESRRG-WT, shRNA directed against ESRRG (shESRRG), or controls for each (GFP and shGFP, respectively). The blot was probed for endogenous total RB1, RB1 phosphorylated at serine 795, the V5 tag, p21, and beta-actin loading control. (**E**) ESRRG ChIP-seq peaks at the CDKN1A locus in RB006 cells (blue) and RB018 cells (green). Corresponding RNA-seq data are shown for RB006 (red), RB016 (green), RB018 (blue), and wRB6 (purple) cells engineered to express shRNA directed against ESRRG (shESRRG) or control (shGFP). P-value was calculated after batch and dispersion correction using EdgeR.

### ESRRG is a hypoxic adaptation and survival factor in Rb cells

Given the known role of ESRRG in hypoxic adaptation in the retina and other tissues (*21, 22*), we investigated how RB1 loss affects regulation of ESRRG under hypoxic conditions. In RB1-wildtype HEK293 cells, hypoxic conditions (1% O2) induced a transient ~3-fold increase in *ESRRG* mRNA expression, which was associated with transient modest activation of the ERRE luciferase reporter (**Fig. 5A**). In HEK293 cells depleted of RB1, *ESRRG* expression levels were over 3-fold higher at baseline and demonstrated an exaggerated increase in response to hypoxia, which was accompanied by hyperactivity of the ERRE reporter over the same time course. Conversely, RB1-null C33A cells demonstrated an exaggerated response to hypoxia in terms of both *ESRRG* expression and ERRE reporter activity, both of which were repressed by ectopic expression of RB1 (**Fig. 5A**). In Rb cell lines, shRNA-mediated depletion of ESRRG or inhibition of ESRRG using the specific inverse agonist GSK5182 (*23*) resulted in marked cell death, which was exacerbated in hypoxia (**Fig. 5B-D**). Taken together, these findings suggest that RB1 dampens the hypoxia-induced surge in ESRRG activity, and that loss of RB1 abolishes this homeostatic mechanism, resulting in nascent Rb cells becoming increasingly dependent on ESRRG in the hypoxic tumor microenvironment. Consistent with this possibility, ESRRG is normally expressed only in isolated retinal cells (**fig. S8**), but it is diffusely expressed in Rb tumors cells, particularly in hypoxic regions beyond ~100 microns of tumor blood vessels (**Fig. 6A**) and in hypoxic vitreous “seeds” (**Fig. 6B**). Intense ESRRG expression was also observed in Rb cells invading the optic nerve (**Fig. 6C**), which is strongly associated with metastasis and poor outcome (*24*).

**Fig. 5.**
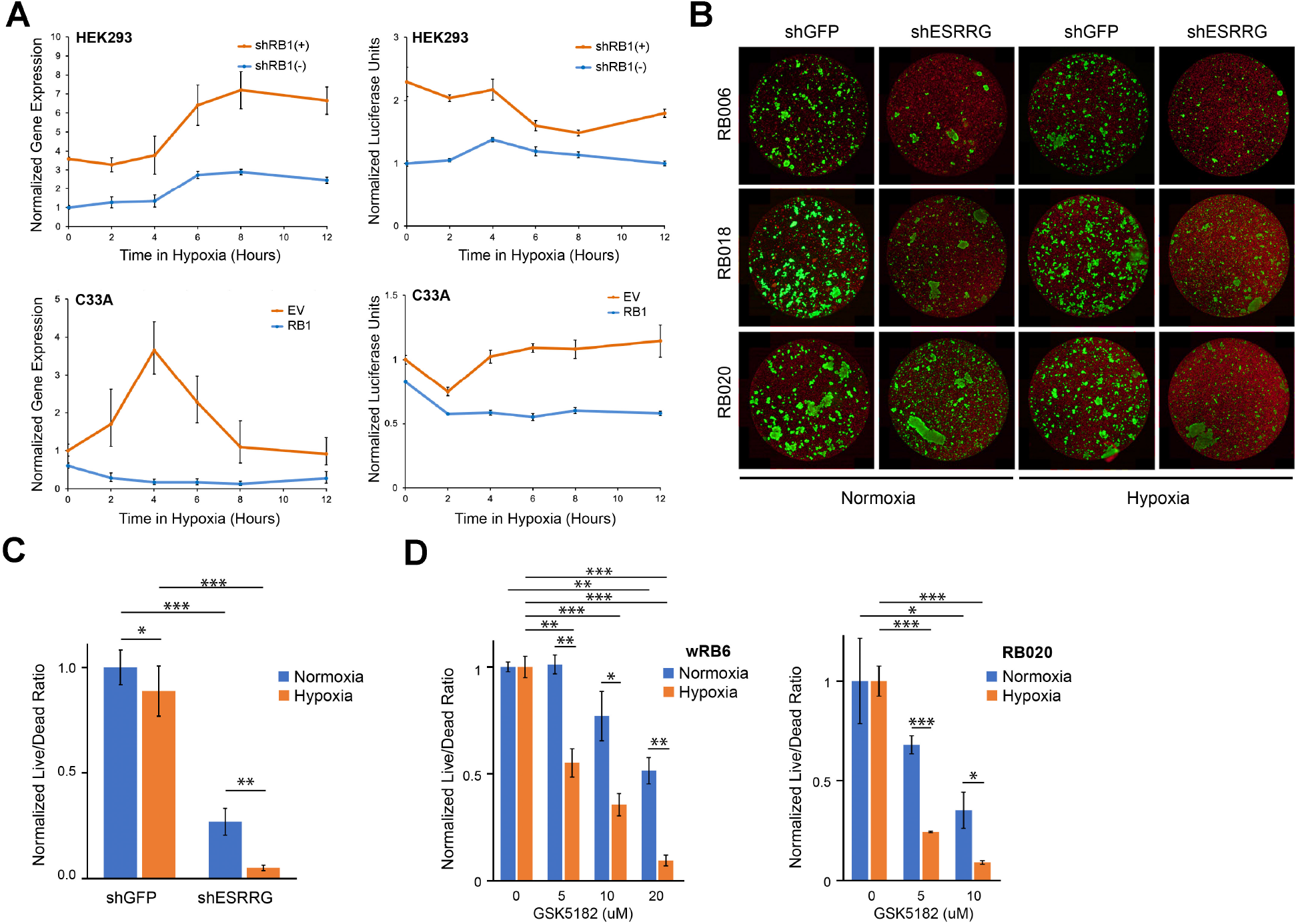
ESRRG is a hypoxic adaptation and survival factor in retinoblastoma. (**A**) Normalized gene expression of ESRRG (upper left) and normalized ERRE luciferase activity (upper right) in HEK293 cells with (DOX+) or without (DOX-) doxycycline treatment to induce shRNA-mediated knockdown of RB1 at the indicated time points in hypoxia (1% O2). Normalized gene expression of ESRRG (lower left) and normalized ERRE luciferase activity (lower right) in C33A cells with ectopically expressed RB1 (RB1) or empty vector control (EV) at the indicated time points in hypoxia (1% O2). (**B**) Representative images of live (green)/dead (red) cell viability assays of 3 recently established Rb cell lines (RB006, RB018, and RB020) stably expressing shRNA directed against ESRRG (shESRRG) or GFP control (shGFP) cultured in normoxia versus hypoxia (1% O2) for 21 days. (**C**) Normalized live/dead cell ratios averaged for all 3 Rb cell lines illustrated in (B) in normoxia versus hypoxia O2 (1%). *, P<0.05; **P<0.01; ***, P<0.001. (**D**) Normalized live/dead cell ratios in wRB6 cells (left) and RB020 cells (right) treated with the indicated micromolar concentrations of the ESRRG inverse agonist GSK5182 for 7 days in normoxia versus hypoxia (1% O2). *, P<0.05; **P<0.01; ***, P<0.001.

**Fig. 6.**
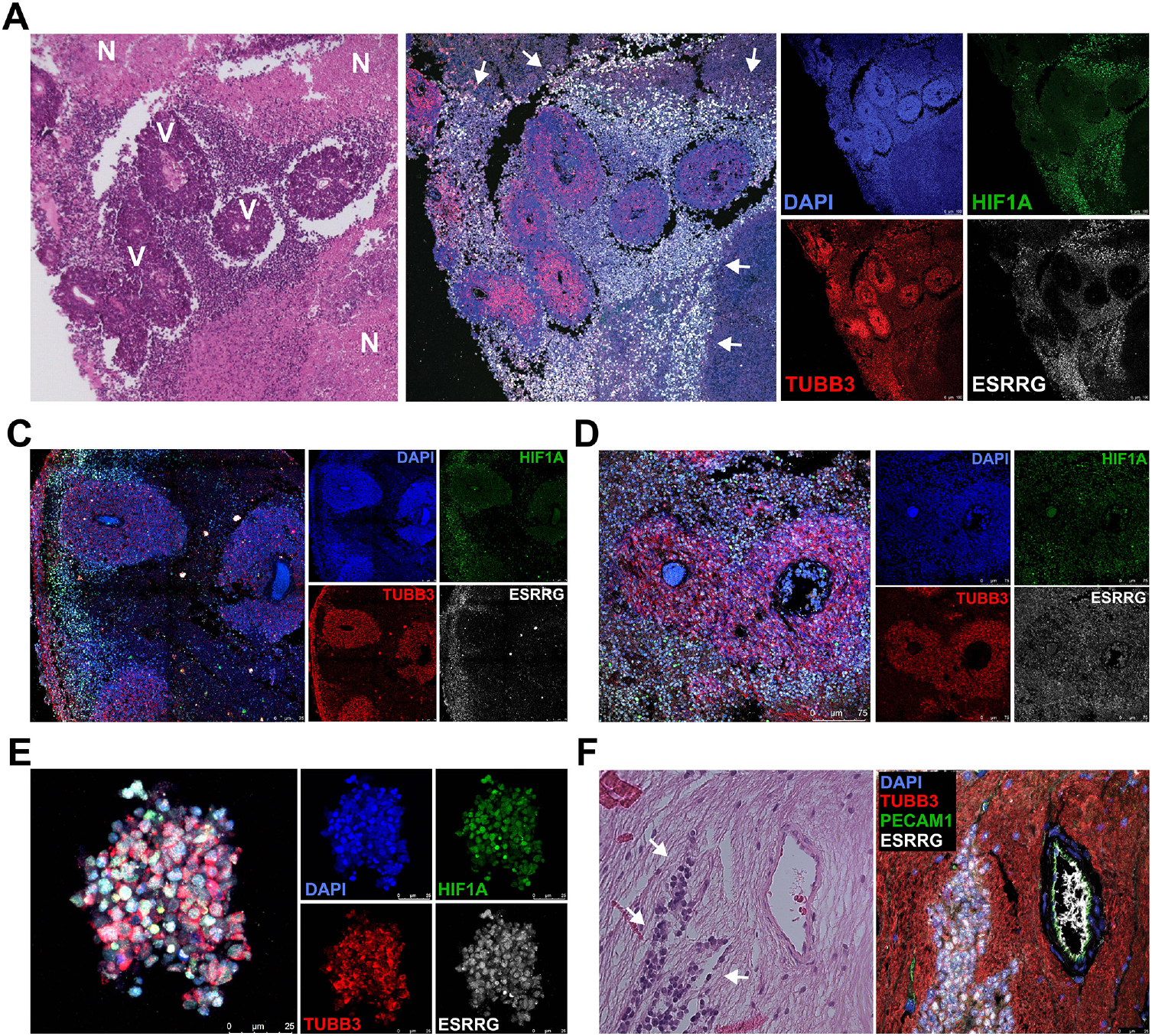
ESRRG expression in human retinoblastomas. (**A**) Hematoxylin and eosin staining (left panel) and multiplex fluorescence immunohistochemistry (middle and right panels) of Rb sample #2587-18. ESRRG (white), TUBB3 (red), DAPI (blue) and HIF1A (green). V, viable tumor cells concentrated in cuffs around blood vessels; N, necrotic tumor cells beyond ~100 microns of tumor blood vessels. Arrows demarcate regions enriched for ESRRG- and HIF1A-expressing cells in hypoxic zones between viable and necrotic tumor cells. (**B-C**) Similar multiplex fluorescence immunohistochemistry findings in Rb samples #2647-18 (left panels) and #1109189582 (right panels). ESRRG (white), TUBB3 (red), DAPI (blue) and HIF1A (green). (**D**) Vitreous tumor cell cluster or “seed” in Rb sample #2647-18 demonstrating strong diffuse staining for ESRRG and HIF1A. ESRRG (white), TUBB3 (red), DAPI (blue) and HIF1A (green). (**E**) Hematoxylin and eosin staining (left panel) and multiplex fluorescence immunohistochemistry (right panel) in Rb sample #29-15 showing invasion of the optic nerve by tumor cells (arrows). ESRRG (white), TUBB3 (red), DAPI (blue) and PECAM1 (green).

## Discussion

In this study, we show that (1) Rb cells are dependent on ESRRG for survival and hypoxic adaptation, (2) RB1 interacts with and inhibits ESRRG such that loss of RB1 releases ESRRG from negative regulation, and (*3*) several recurring genomic aberrations in Rb increase ESRRG dosage and/or activity. The most common CNV in Rb is gain of chromosome 1q, which occurs in two-thirds of cases and increases the gene dosage of *ESRRG*, located at 1q41 (**Fig. 1**)(*7, 10*). Although several genes on chromosome 1q have been postulated as possible drivers of Rb progression (*5, 25*), these claims have not been functionally validated. Our functional evidence raises the interesting possibility that ESRRG may be the main driver of Rb progression on 1q that accounts for the frequent gain of this chromosomal segment. After *RB1*, the most commonly mutated gene in Rb is *BCOR*, a transcriptional co-repressor and component of the non-canonical PRC1.1 polycomb complex that is also mutated in other developmental disorders affecting the eye (*26, 27*). Inactivating mutations in *BCOR* are found in about 20% of Rb tumors and are associated with poor prognosis (*10, 28*). We show here that BCOR represses ERRE-mediated transcription independently of RB1, suggesting that BCOR loss derepresses ESRRG beyond that associated with RB1 loss. The pattern of recurrent genomic aberrations in Rb is consistent with ongoing selective pressure to increase ESRRG activity during tumor evolution, which could make ESRRG an attractive therapeutic target.

ESRRG is an estrogen-related orphan nuclear receptor transcription factor that is normally expressed in the developing retina and central nervous system, where it regulates genes involved in development, proliferation, and oxygen metabolism (*21, 22, 29–31*). In Rb cells, we found that ESRRG localizes to and regulates the transcription of numerous genes containing CRX, OTX2, NEUROD, and LHX binding motifs that regulate normal retinal development (*32–34*). CRX and OTX2 in particular are key regulators of oxidative metabolism and mitochondrial function during early neurogenesis and retinogenesis (*35–37*), and they can become oncogenic when aberrantly activated (*38, 39*). The developing retina is a hypoxic environment, owing to the proliferation of retinal precursor cells, oxygen demand of newly formed neurons, and limited blood supply (*40, 41*). This oxygen demand is markedly increased in Rb due to rapid uncontrolled proliferation of tumor cells (*42, 43*). We found that Rb cells require ESRRG for survival, proliferation, and hypoxic adaptation. Indeed, Rb cells demonstrate constitutive ESRRG expression that is enriched in oxygen-poor regions beyond ~100 microns from tumor blood vessels (**Fig. 6A**) (*44*) and in vitreous “seeds” (**Fig. 6B**), which consist of clusters of Rb cells that proliferate in the hypoxic vitreous cavity and portend chemoresistance and poor outcome in Rb (*45, 46*). Further, ESRRG expression clearly demarcates aggressive Rb cells invading the optic nerve (**Fig. 6C**), which is a major risk factor for metastatic death(*24*).

In conclusion, Rb tumors undergo ongoing hypoxic adaptation to allow continued proliferation and invasion in an increasingly oxygen-depleted environment (*47, 48*), and our findings implicate ESRRG and its decoupling from normal homeostatic regulation by RB1 and BCOR as a key mediator of this hypoxic adaptation. Further studies are warranted to investigate the potential role for inhibiting ESRRG in the management of Rb.

## Materials and Methods

### Specimens and clinical data

Human Rb samples were collected from enucleated eyes at the time of surgery, snap frozen, and stored at −80C until used for analysis with the approval of the University of Miami Institutional Review Board. Clinical and histopathologic information were obtained and samples de-identified for further analysis.

### DNA sequencing, quality control, and alignment

WES was performed on 31 of our Rb enucleated tumor samples, 20 of which had matched blood DNA available for sequencing. DNA was extracted using the Wizard Genomic DNA Purification kit (Promega, Madison, WI) and the Quick Gene DNA whole blood kit S (Fugifilm, Tokyo, Japan), respectively. Exome fragments were captured using the SureSelect Human Exon V5 (Agilent) and sequenced on the Illumina HiSeq 2500. WES data from an additional 72 Rb samples with matched blood was obtained from the European Genome-Phenome Archive (EGAS00001001690) with permission from the Data Access Committee (EGAC00001000431)(*7*). WGS from 4 Rb samples with matched blood was obtained with permission from dbGaP (phs000352.v1.p1). Fastq files from all sources underwent the following bioinformatics pipeline. Sequence data were quality controlled using FastQC (v0.11.3). WES and WGS reads were trimmed (if required) and aligned to the human genome (hg19/GRCH37) using Novoalign (v3.06.04), marked for duplicates using Picard (v1.128), realigned around small and large indels using Abra (v0.94c)(*49*), and read mate fixed and analyzed for coverage statistics using Picard.

### Mutation analysis

WES datasets underwent variant calling for SNPs and Indels using MuTect2 (GATK 2016-01-25 nightly build) (*50*). For MuTect2, mutation calls for a panel of normals (n=117 germline blood samples) was generated and pooled together to further filter out mutations that were present in at least two normals. For tumors without matched blood samples, MuTect2 was used for variant calling with a panel of normals and a high coverage blood sample to filter out likely germline mutations. Mutations were further filtered out if both alternate tumor read counts were < 3, if alternate tumor read counts were <8, and blood alternate read counts were >1, and the minor allele frequency (MAF) was < 5%. For all sequencing samples, the BAM files were investigated manually for the regions of interest (RB1, BCOR) using the Integrative Genomics Viewer (IGV)(v2.3.80, Broad Institute, Cambridge, MA).

For all called mutations, ANNOVAR was used for annotation (*51*). Following annotation, mutations were filtered out if the MAF was 1% or greater in the population according to 1000 Genomes Project (August 2015) or Exome Sequencing Project (2015 Mar), and mutations listed in dbSNP (v138) were filtered out, with the exception of SNPs with a MAF < 1% (or unknown) in the population, a single mapping to reference assembly, or with a “clinically associated” tag. If the exact same insertion or deletion or non-frameshift multi-nucleotide substitution was present in at least 2 samples without matched germline and were not present in any matched tumor sample, it was removed as a likely germline variant. Functional consequences of SNPs were assessed by three mutational predictor tools: Polyphen2 (probably damaging, possibly damaging, benign) (*52*), FATHMM (damaging, tolerated) (*53*), and MetaLR (damaging, tolerated) (*54*). In non exonic regions and for synonymous mutations, SNPs were considered deleterious if two out of three of the above prediction algorithms predicted a damaging mutation. For non-synonymous exonic SNPs, mutations were not considered deleterious if 2 out of 3 algorithms predicted a benign or tolerated mutation. Probably damaging, possibly damaging, or damaging calls were considered deleterious. All insertions and deletions in exonic regions and alterations in splicing junctions were considered deleterious. For identification of additional driver mutations, the unmatched tumor samples were first excluded before being evaluated if mutations were found in matched samples and were also excluded if known to be one of the benign mutations commonly seen in cancer exomes (*55, 56*). Lollipop plots displaying the distribution of driver mutations along the protein domains of each gene were plotted using trackViewer (*57*) in R (version 4.0.3).

### Copy number variations

CNVs were ascertained from WES data using CNVKit (v0.8.5) (*58*) and by cgpBattenberg using default settings (*59*). Isodisomy was determined by plotting B allele frequencies (BAF) using CNVKit, ASCAT, and cgpBattenberg. CNVKit output was analyzed using GISTIC (*60*) to determine significant CNVs using cutoffs of FDR < 0.05, Z-score ≥ 6, and chromosomal arm score > |0.1|. MYCN amplification was determined using GISTIC with a Z-score cutoff of 3. Structural variants from the WGS samples were determined using DELLY (version 0.7.6) (*61*).

### Pathway analysis

Genes with called mutations were analyzed for functional relationships with the Ingenuity Pathway Analysis product (QIAGEN, Valencia, CA, USA), using the Path Explorer option of the Path Designer tool. The analysis was filtered for paths with a maximum of 2 nodes between selected genes.

### Cell culture

Newly established, patient-derived low passage Rb cell lines (wRB6, RB006, RB010, RB015, RB016, RB018, RB020, and RB021) were derived from enucleated eyes of patients of the senior author (J.W.H.) and cultured at 5% O2 in Dulbecco’s modified essential medium (DMEM)/F12 with B-27 minus vitamin A (Life Technologies), 1% penicillin-streptomycin, 2 mM Glutamax (Gibco), 10 ng/mL basic fibroblast growth factor (bFGF)(PeproTech, catalogue #100-18B), 10 ng/mL recombinant human stem cell factor (rSCF)(PeproTech, catalogue #300-07), and 20 ng/mL epidermal growth factor (EGF)(PeproTech AF-100-15). C33A cells were obtained from ATCC and maintained in Eagle’s Minimum Essential Medium supplemented with 10% heat-inactivated fetal bovine serum (HI-FBS) and 1% penicillin-streptomycin. HEK293 cells were obtained from ATCC and maintained in DMEM media with 10% HI-FBS, 2mM Glutamax, and 1% penicillinstreptomycin. Cell proliferation and viability were measured using Trypan Blue Stain (Gibco) and/or LIVE/DEAD Cell Viability Assay (ThermoFischer Scientific). All cells were maintained in ambient O2 unless otherwise stated. Rb cells were treated with various concentrations of the ESRRG selective inverse agonist GSK5182 (Aobious, Inc., Gloucester, MA) for 7 days.

### Plasmids and lentivirus expression vectors

Short hairpin RNA (shRNA) plasmids were created by integrating validated siRNA sequences CCTGTCAGGAAACTGTATGAT (*62*) and AATGGCCATCAGAACGGACTT targeting human ESRRG (shESRRG) into pLKO.1-Puro vector (Addgene #8453). Non-specific siRNA sequence AACAGCCACAACGTCTATATC or siRNA sequence CAACAGCCACAACGTCTATAT targeting GFP were used as controls for shRNA. Tetracycline-inducible TET-shRB1 and constitutive shRB1 constructs were created by integrating validated siRNA sequences GAAAGGACATGTGAACTTA (*63*) and GAACGATTATCCATTCAAA (*64*) targeting human RB1 (shRB1) into TET-pLKO.1-Puro vector (Addgene #21915) and pLKO.1-Puro vector, respectively. The plasmids were packaged into lentiviral particles by transient co-transfection into HEK293T cells with pMD2G and psPAX2 packaging plasmids using JetPrime reagent (Polyplus). The pCMV-ESRRG-WT-V5 vector was created by PCR amplification of ESRRG full length cDNA fragment (Horizon #MHS6278-202806209) and subsequent recombination into a pCMV plasmid encoding C-terminal V5. An RB1-binding site mutant of ESRRG (V5-ESRRG-MUT) was generated using site-directed mutagenesis to replace 5’-GTCAGGAAACTGTATGATGAC-3’ with 5’-GACAAAGTCGATGCGAGGTAT-3’. A construct encoding RB1 was created by PCR-amplifying cDNA for human RB1(Horizon #MHS6278-202758572) and cloning into an N-terminal HA-tagged expression vector. The pLL-FLAG-BCOR plasmid was a gift from Dr. Vivian Bardwell. The ERRE promoter luciferase reporter plasmid 3xERRE-luciferase (pERRE-LUC; Addgene # 37851) was a gift from Dr. Rebecca Riggins. CMV-promoter-driven beta-galactosidase plasmid (pCMV-beta-gal) was obtained from ATCC (#77177). Lentiviral particles were concentrated by precipitation from culture supernatants using PEG-IT (System Biosciences) and quantified with the QuickTiter Lentivirus Titer Kit (Cell Bio Labs).

### Luciferase assays

The RB1-null cell line C33A was co-transfected with indicated quantities of pCMV-HA-RB1, pLL-FLAG-BCOR, pCMV-V5-ESRRG-WT or pCMV-V5-ESRRG-MUT, and pERRE-LUC using JetPrime reagent (Polyplus). Stoichiometric quantities of empty vectors pCMV-V5-EV, pCMV-HA-EV and pLL-FLAG-EV were used for C33A cell transfection normalization. H293 cell expressing TET-shRB1 (H293-TET-shRB1) were induced with doxycycline for 48 hrs then co-transfected with indicated quantities pCMV-HA-RB1, pCMV-V5-ESRRG-WT or pCMV-V5-ESRRG-MUT, and pERRE-LUC. Cells not treated with doxycycline and stoichiometric quantities of empty vectors pCMV-V5-EV and pLL-FLAG-EV were used for H293 cell transfection normalization. Luciferase activity was measured 24 hours after plasmids transduction by lysing the cells in the Passive Buffer (Promega) and combining equal volumes of cell lysates with the Bright-Glo™ Reagent (Promega). Co-transfection with a CMV-beta-galactosidase plasmid and bicinchoninic acid protein quantification assays were used for luciferase signal normalization.

### Co-immunoprecipitations

RB1-null C33A cells (10^7^ cells per condition) were cultured in 15 cm dishes, transfected with the indicated expression plasmids, and lysed with ice-cold NP-40 buffer supplemented with protease and phosphatase inhibitors (Roche). Cell lysates were passed 20 times through a 25-gauge needle, incubated on ice for 1 hour, pelleted by centrifugation at 12000 RCF for 10 min, precleared by 30 min co-incubation with magnetic beads (Life Technologies) at 4°C, and incubated overnight at 4°C with 5 micrograms of the indicated antibodies per 1 mg of quantified protein lysates. Antibody-protein conjugates were captured by incubation with protein-G Dynabeads for 1 hour, washed with ice-cold Co-IP buffer, and eluted by boiling in SDS-running buffer. Total precleared lysates were used as positive control.

### Western blot

Cell pellets were resuspended in lysis buffer, which consisted of 50mM Hepes pH7.2, 400mM NaCl, 0.1% NP-40, 0.5mM EDTA pH8, 2.5mM DTT, plus protease and phosphatase inhibitors. Resulting cell lysates were incubated on ice for 10 mins, sonicated at medium power for 10 sec, centrifuged for 10 min at 10000 RCF to remove cellular debris, quantified, and equal quantities denatured by boiling with SDS loading buffer for 5 minutes at 90°C. Proteins (30 micrograms per condition) were resolved on SDS-PAGE gradient gels and transferred to a polyvinylidene difluoride (PVDF) membrane (ThermoFisher Scientific). Blots were blocked in 5% bovine serum albumin (BSA) in Tris-buffered saline (TBS; pH 7.6); probed overnight with indicated primary antibodies in 5% BSA and 0.15% Tween20 in TBS (TBS-T), washed in TBS-T, and incubated for 1 hour with a secondary antibody in 5% BSA/TBS-T solution. Proteins were visualized using SuperSignal chemiluminescence (ThermoFisher Scientific) or LI-COR imaging technology (LI-COR Biosciences).

### Hypoxia experiments

Recently established Rb cell lines (RB006, RB018, RB020, and RB021) were transduced with either shESRRG- or shCTRL lentiviral particles and treated with 2 micrograms per mL puromycin 48 hrs after transduction to select for stable lentiviral integration. At 7 days after transduction, cells were maintained in normoxia or hypoxia (1% O2). Cell viability at day 21 was measured with the LIVE/DEAD cell viability assay (ThermoFisher Scientific). Whole wells were scanned using the EVOS FL Auto Imaging System (ThermoFisher Scientific), and image fluorescence was quantified using particle analysis with ImageJ software. HEK293 stably integrated with TET-shRB1 were induced with 1 microgram per mL doxycycline hyclate for 48 hrs, transfected with pERRE-LUC plasmid and maintained in either normoxia or hypoxia (1% O2) for an additional 24 hours. C33A cells were transfected with pCMV-HA-RB1 and pERRE-LUC plasmids for 24 hrs and exposed to either normoxia or hypoxia (1% O2) for 24 hours. Uninduced H293-TET-shRB1 and C33A cells transfected with pCMV-HA-EV were used as experimental controls.

### Quantitative real-time polymerase chain reaction

Total RNA was extracted from cells with a combination of TriZol and RNeasy Mini RNA isolation kit (QIAGEN, Valencia, CA, USA), and reverse-transcribed using high capacity cDNA transcription kit (Applied Biosystems, Foster City, CA, USA). Quantitative polymerase chain reaction (qPCR) was performed in triplicate using the 7300 Real-time RT-PCR system (Applied Biosystems) according to the manufacturer’s description using the following thermocycler parameters: 1 minute of initial activation at 98°C, 40 cycles of 5 seconds denaturation at 98°C, and 55 seconds annealing and extension at 60°C. The relative gene expression data were analyzed by the comparative CT (ΔΔCT) method. The results were normalized to GAPDH, RNA18S, and ACTB as internal controls.

### Chromatin immunoprecipitation and DNA sequencing (ChIP-seq)

Recently established RB006 and RB018 cells were grown to approximately 5×10^7^, collected and pelleted. Cell pellets were fixed in 1% formaldehyde for 10 minutes to crosslink protein-DNA interactions. Cell nuclei were extracted and subjected to sonication in a Biorupter Pico (Diagenode). Shearing was optimized to yield chromatin fragments of 200-500bp. Chromatin was incubated in ChIP-qualified antibody against ESRRG. Antibodies were captured using protein-G coupled magnetic beads and protein-DNA was eluted. DNA was purified for downstream analysis. Successful ChIP was validated with RT-PCR for known binding genes and known negative controls. Technical controls included non-specific IgG incubation of cell lysates as well as no-antibody (input) controls for non-specific DNA binding to protein-G coupled beads. DNA was sent for library preparation and sequencing in the Oncogenomics Shared Resource of the University of Miami. Bioinformatic analysis was performed by established ChIP-seq pipelines including quality control (FastQC), adapter trimming and alignment (Novoalign), and normalization to an input control followed by peak calling (MACS2). Heatmap and bandplots were generated using Deeptools. Overlapping high-confidence peaks that were called in both the RB006 and RB018 ESRRG-pulldown ChIP-Seq datasets as well as a pooled dataset that combined the RB006 and RB018 data were determined using HOMER (p<0.001) and used for downstream analyses. Euler diagram of peaks called for each dataset and overlapping datasets was plotted using EulerAPE (version 3.0) (*65*). Motif and gene set enrichment analyses were conducted using HOMER. Super enhancer regions identified in retinal and neural tissue were downloaded from the comprehensive human Super-Enhancer database (SEdb, version 1.03)(*14*). Peaks were plotted for visualization using our custom program SparK (*66*) (https://github.com/HarbourLab/SparK). Peak location, motif analysis, and co-occurrence visualization were conducted using ChIPseeker, UpSetR, and custom scripts (available upon request)(*67, 68*).

### RNA sequencing (RNA-seq) and analysis

RNA was extracted from tumor samples or Rb cell lines (RB006, RB016, RB018, and wRB6) with or without shESRRG knockdown using a combination of TriZol and the RNeasy Mini RNA isolation kit (QIAGEN, Valencia, CA, USA). RNA-seq libraries were prepared using the TruSeq Stranded Total RNA prep kit with Ribo-Zero Gold to remove cytoplasmic and mitochondrial rRNA according to the manufacturer’s recommendation (Illumina, San Diego, CA). Total RNA-seq libraries were run on an Illumina NextSeq 500 sequencing instrument according to the manufacturer’s protocol. Reads were aligned and gene counts were made using STAR, data quality was assessed using FastQC and RSeQC, gene expression was normalized, batch corrected, and determined using EdgeR. Significant differences (FDR<0.05) between cells expressing shESRRG versus shGFP control were calculated using EdgeR. For tumor samples that underwent RNA sequencing without a detectable biallelic loss of RB1 seen in exome analysis, RNA tracks were manually curated to look for any *RB1* mutations.

### Integration of ChIP-seq and RNA-seq data

Significant ESRRG ChIP-Seq peaks overlapping between RB006, RB018, and pooled samples were correlated with significantly up-regulated or down-regulated genes after shESRRG knockdown in 4 primary-cultured cell lines (RB006, RB016, RB018, and wRB6). Genes that were significantly up-regulated or down-regulated in association with ESRRG ChIP-Seq peaks were analyzed with GSEA and MSigDB. Differentially expressed genes after ESRRG knockdown were evaluated for ESRRG-associated transcription factor motifs.

### Single-cell RNA-seq analysis

Single-cell suspensions were counted using the Cellometer K2 Fluorescent Viability Cell Counter (Nexcelom), verified manually using a hemocytometer, and adjusted to 1,000 cells/ul. For RNA sequencing, samples were prepared using the Chromium Single Cell 5’ Library & Gel Bead Kit v2 (10X Genomics) according to the manufacturer’s protocol with a capture target of 10,000 cells. Each sample was processed on an independent Chromium Single Cell A Chip (10X Genomics) and run on a thermocycler (Eppendorf). 5’ gene expression libraries were sequenced using NextSeq 500 150-cycle high-output flow cells or NovaSeq 6000 200 cycle S4 flow cells. 5’ Single-cell RNA-Seq Analysis. Raw base call (BCL) files were analyzed using CellRanger (version 3.0.2). The “mkfastq” command was used to generate FASTQ files and the “count” command was used to generate raw gene-barcode matrices aligned to the GRCh38 Ensembl build 93 genome. The data from all five samples were combined in R (3.5.2) using the Seurat package (3.0.0) and an aggregate Seurat object was generated (*69, 70*). To ensure our analysis was on high-quality cells, filtering was conducted by retaining cells that had unique molecular identifiers (UMIs) greater than 400, expressed 100 to 6000 genes inclusive, and had mitochondrial content less than 10 percent. This resulted in 6,371 cells for RB025, 12,808 cells for RB026, 10,675 cells for RB027, 1,354 cells for RB028, and 1,445 cells for RB029. Doublets were assessed using the DoubletFinder (2.0.3) algorithm(*71*) using the multiplet rates in the 10X Genomics Chromium Single Cell 3’ Reagent Kits User Guide (v3 Chemistry). Doublets were assessed by individual samples based on the number of captured cells. Data for all 5 samples were combined using the Standard Integration Workflow (https://satijalab.org/seurat/v3.0/integration.html). Data from each sample were normalized using the NormalizeData() function and variable features were identified using FindVariableFeatures() with 5000 genes and the selection method set to “vst”, a variance stabilizing transformation. To identify integration anchor genes among the five samples the FindIntegrationAnchors() function was used with 30 principal components and 5000 genes. Using Seurat’s IntegrateData() the samples were combined into one object. The data were scaled using the ScaleData() function to reduce dimensionality of this dataset. Principal component analysis (PCA) was performed, and the first 30 principal components were summarized using uniform manifold approximation and projection (UMAP) dimensionality reduction(*72*). 30 principal components (PCs) were chosen based on results from analysis using JackStraw() and elbow plots. The DimPlot() function was used to generate the UMAP plots (**Fig. 2C**). Clustering was conducted using the FindNeighbors() and FindClusters() functions using 30 PCA components and a resolution parameter set to 1.8. The original Louvain algorithm was used for modularity optimization(*73*). The resulting 36 louvain clusters were visualized in a two-dimensional UMAP representation. Differentially expressed genes were identified for each cluster using FindAllMarkers() and filtered for |log fold-change| greater than 0.5 and adjusted p-value less than 0.05. Dot plots were generated using the DotPlot() function, Violin plots were generated with the vlnplot() function, and scatter plots were generated with FeatureScatter() in Seurat with “assay” set to “RNA”.

### Immunohistochemistry

For immunohistochemistry, 4-μm paraffin sections of Rb tumor specimens were de-paraffinized in xylene, rehydrated in an ethanol gradient, permeabilized for 30 minutes in 0.5% Triton X-100 (v/v in PBS), and subjected to antigen retrieval using sodium citrate buffer (pH 6.0) at 95°C for 20 minutes. Sections were then washed, equilibrated in water for 1 hour, and blocked with 1% BSA for 1 hour at room temperature. Samples were then incubated in primary antibody overnight at 4°C, washed 3x in distilled water and then incubated in secondary antibody for 2 hours at room temperature. Samples were then washed 3x in distilled water, mounted with SlowFade Diamond Antifade mounting medium with DAPI (ThermoFisher Scientific), and imaged using an SP8 Leica laser scanning confocal microscope (Leica Microsystems Inc, Buffalo Grove, IL).

### Statistical analyses

Data statistics and bioinformatics analyses were performed using R (version 4.0.3) and Bioconductor packages (https://www.bioconductor.org/) unless otherwise indicated. Multiple comparisons were adjusted by the Benjamini-Hochberg correction.

## Acknowledgments

We acknowledge the support of the Sylvester Comprehensive Cancer Center Biostatistics & Bioinformatics Shared Resource (BBSR) and Oncogenomics Shared Resource (OGSR), and the University of Miami Institute for Data Science and Computing (IDSC).

## Funding

Alex’s Lemonade Stand/Tap Cancer Out Innovation Grant (JWH, DP)

University of Miami Sheila and David Fuente Graduate Program in Cancer Biology (MGF, MAD, MGZ, JJD)

University of Miami Center for Computational Science Fellowship (MGF, MAD)

University of Miami Lois Pope LIFE Fellowship Award (MGF)

Alcon Research Award (JWH)

NIH/NCI R01CA248890 (DP, JWH)

NIH/NCI P30CA240139 (Sylvester Comprehensive Cancer Center)

NIH/NEI P30EY014801 (Bascom Palmer Eye Institute)

Research to Prevent Blindness Unrestricted Grant (Bascom Palmer Eye Institute)

Generous philanthropic gift from Dr. Mark J. Daily (JWH)

The content is solely the responsibility of the authors and does not necessarily represent the official views of the National Institutes of Health.

## Author contributions

Conceptualization: MGF, DP, JWH

Methodology: MGF, JNK, DP, JWH

Software: MGZ, JJD, MAD, SK

Investigation: MGF, JNK, MGZ, YS, JW, ZAT

Data Curation: CLD

Supervision: DP, JWH

Writing—original draft: MGF, JWH

Writing—review & editing: JNK, MGZ, JJD, MAD, CLD, SK, DP

## References

1. I. D. Fabian et al., Global Retinoblastoma Presentation and Analysis by National Income Level. JAMA Oncol 6, 1–12 (2020).

2. H. E. Grossniklaus, Retinoblastoma. Fifty years of progress. The LXXI Edward Jackson Memorial Lecture. Am J Ophthalmol 158, 875–891 (2014).

3. S. H. Friend et al., A human DNA segment with properties of the gene that predisposes to retinoblastoma and osteosarcoma. Nature 323, 643–646 (1986).

4. X. L. Xu et al., Rb suppresses human cone-precursor-derived retinoblastoma tumours. Nature 514, 385–388 (2014).

5. T. W. Corson, B. L. Gallie, One hit, two hits, three hits, more? Genomic changes in the development of retinoblastoma. Genes Chromosomes Cancer 46, 617–634 (2007).

6. H. Dimaras et al., Loss of RB1 induces non-proliferative retinoma: increasing genomic instability correlates with progression to retinoblastoma. Hum Mol Genet 17, 1363–1372 (2008).

7. I. E. Kooi et al., Somatic genomic alterations in retinoblastoma beyond RB1 are rare and limited to copy number changes. Sci Rep 6, 25264 (2016).

8. K. Sampieri et al., Array comparative genomic hybridization in retinoma and retinoblastoma tissues. Cancer Sci 100, 465–471 (2009).

9. N. A. Laurie et al., Inactivation of the p53 pathway in retinoblastoma. Nature 444, 61–66 (2006).

10. J. H. Francis et al., Molecular Changes in Retinoblastoma beyond RB1: Findings from Next-Generation Sequencing. Cancers (Basel) 13, (2021).

11. H. R. Davies et al., Whole-Genome Sequencing of Retinoblastoma Reveals the Diversity of Rearrangements Disrupting RB1 and Uncovers a Treatment-Related Mutational Signature. Cancers (Basel) 13, (2021).

12. A. R. Afshar et al., Next-generation sequencing of retinoblastoma Identifies pathogenic alterations beyond RB1 inactivation that correlate with aggressive histopathologic features. Ophthalmology, (2019).

13. A. Kramer, J. Green, J. Pollard, Jr., S. Tugendreich, Causal analysis approaches in Ingenuity Pathway Analysis. Bioinformatics (Oxford, England) 30, 523–530 (2014).

14. Y. Jiang et al., SEdb: a comprehensive human super-enhancer database. Nucleic acids research 47, D235–D243 (2019).

15. B. Xiao et al., Crystal structure of the retinoblastoma tumor suppressor protein bound to E2F and the molecular basis of its regulation. Proc Natl Acad Sci U S A 100, 2363–2368 (2003).

16. C. Lee, J. H. Chang, H. S. Lee, Y. Cho, Structural basis for the recognition of the E2F transactivation domain by the retinoblastoma tumor suppressor. Genes Dev 16, 3199–3212. (2002).

17. W. D. Cress, D. G. Johnson, J. R. Nevins, A genetic analysis of the E2F1 gene distinguishes regulation by Rb, p107, and adenovirus E4. Molecular and cellular biology 13, 6314–6325 (1993).

18. H. Dinkel et al., ELM 2016--data update and new functionality of the eukaryotic linear motif resource. Nucleic acids research 44, D294–300 (2016).

19. R. Sladek, J. A. Bader, V. Giguere, The orphan nuclear receptor estrogen-related receptor alpha is a transcriptional regulator of the human medium-chain acyl coenzyme A dehydrogenase gene. Molecular and cellular biology 17, 5400–5409 (1997).

20. S. Yu, X. Wang, C. F. Ng, S. Chen, F. L. Chan, ERRgamma suppresses cell proliferation and tumor growth of androgen-sensitive and androgen-insensitive prostate cancer cells and its implication as a therapeutic target for prostate cancer. Cancer Res 67, 4904–4914 (2007).

21. J. Y. Do et al., Retinal hypoxia induces vascular endothelial growth factor through induction of estrogen-related receptor gamma. Biochem Biophys Res Commun 460, 457–463 (2015).

22. P. Kumar, C. R. Mendelson, Estrogen-related receptor gamma (ERRgamma) mediates oxygen-dependent induction of aromatase (CYP19) gene expression during human trophoblast differentiation. Mol Endocrinol 25, 1513–1526 (2011).

23. E. Y. Chao et al., Structure-guided synthesis of tamoxifen analogs with improved selectivity for the orphan ERRgamma. Bioorg Med Chem Lett 16, 821–824 (2006).

24. P. T. Finger, J. W. Harbour, Z. A. Karcioglu, Risk factors for metastasis in retinoblastoma. Survey of ophthalmology 47, 1–16 (2002).

25. I. E. Kooi et al., A Meta-Analysis of Retinoblastoma Copy Numbers Refines the List of Possible Driver Genes Involved in Tumor Progression. PLoS One 11, e0153323 (2016).

26. J. A. Simon, R. E. Kingston, Mechanisms of polycomb gene silencing: knowns and unknowns. Nat Rev Mol Cell Biol 10, 697–708 (2009).

27. D. Ng et al., Oculofaciocardiodental and Lenz microphthalmia syndromes result from distinct classes of mutations in BCOR. Nature genetics 36, 411–416 (2004).

28. Y. Yamamoto, A. Abe, N. Emi, Clarifying the impact of polycomb complex component disruption in human cancers. Mol Cancer Res 12, 479–484 (2014).

29. I. Hermans-Borgmeyer, U. Susens, U. Borgmeyer, Developmental expression of the estrogen receptor-related receptor gamma in the nervous system during mouse embryogenesis. Mech Dev 97, 197–199 (2000).

30. L. Pei et al., Dependence of hippocampal function on ERRgamma-regulated mitochondrial metabolism. Cell Metab 21, 628–636 (2015).

31. A. Ao, H. Wang, S. Kamarajugadda, J. Lu, Involvement of estrogen-related receptors in transcriptional response to hypoxia and growth of solid tumors. Proc Natl Acad Sci U S A 105, 7821–7826 (2008).

32. P. A. Ruzycki, X. Zhang, S. Chen, CRX directs photoreceptor differentiation by accelerating chromatin remodeling at specific target sites. Epigenetics Chromatin 11, 42 (2018).

33. T. J. Cherry et al., NeuroD factors regulate cell fate and neurite stratification in the developing retina. J Neurosci 31, 7365–7379 (2011).

34. R. Balasubramanian, A. Bui, Q. Ding, L. Gan, Expression of LIM-homeodomain transcription factors in the developing and mature mouse retina. Gene Expr Patterns 14, 1–8 (2014).

35. H. T. Kim et al., Mitochondrial Protection by Exogenous Otx2 in Mouse Retinal Neurons. Cell Rep 13, 990–1002 (2015).

36. H. Yamamoto, T. Kon, Y. Omori, T. Furukawa, Functional and Evolutionary Diversification of Otx2 and Crx in Vertebrate Retinal Photoreceptor and Bipolar Cell Development. Cell Rep 30, 658–671 e655 (2020).

37. Y. Satou et al., Phosphorylation states change Otx2 activity for cell proliferation and patterning in the Xenopus embryo. Development 145, (2018).

38. J. Li et al., OTX2 is a therapeutic target for retinoblastoma and may function as a common factor between C-MYC, CRX, and phosphorylated RB pathways. Int J Oncol 47, 1703–1710 (2015).

39. D. D. Glubrecht, J. H. Kim, L. Russell, J. S. Bamforth, R. Godbout, Differential CRX and OTX2 expression in human retina and retinoblastoma. J Neurochem 111, 250–263 (2009).

40. J. S. Joyal, M. L. Gantner, L. E. H. Smith, Retinal energy demands control vascular supply of the retina in development and disease: The role of neuronal lipid and glucose metabolism. Prog Retin Eye Res 64, 131–156 (2018).

41. N. D. Wangsa-Wirawan, R. A. Linsenmeier, Retinal oxygen: fundamental and clinical aspects. Archives of ophthalmology (Chicago, Ill.: 1960) 121, 547–557 (2003).

42. Q. Yang, A. Tripathy, W. Yu, C. G. Eberhart, L. Asnaghi, Hypoxia inhibits growth, proliferation, and increases response to chemotherapy in retinoblastoma cells. Exp Eye Res 162, 48–61 (2017).

43. J. Sudhakar et al., Hypoxic tumor microenvironment in advanced retinoblastoma. Pediatr Blood Cancer 60, 1598–1601 (2013).

44. M. N. Burnier, I. W. McLean, L. E. Zimmerman, S. H. Rosenberg, Retinoblastoma. The relationship of proliferating cells to blood vessels. Investigative ophthalmology & visual science 31, 2037–2040 (1990).

45. U. Winter et al., Tridimensional Retinoblastoma Cultures as Vitreous Seeds Models for Live-Cell Imaging of Chemotherapy Penetration. Int J Mol Sci 20, (2019).

46. K. Gunduz et al., Causes of chemoreduction failure in retinoblastoma and analysis of associated factors leading to eventual treatment with external beam radiotherapy and enucleation. Ophthalmology 111, 1917–1924 (2004).

47. B. F. Fernandes et al., Hypoxia-inducible factor-1alpha and its role in the proliferation of retinoblastoma cells. Pathol Oncol Res 20, 557–563 (2014).

48. Y. Y. Li, Y. L. Zheng, Hypoxia promotes invasion of retinoblastoma cells in vitro by upregulating HIF-1α/MMP9 signaling pathway. Eur Rev Med Pharmacol Sci 21, 5361–5369 (2017).

49. L. E. Mose, M. D. Wilkerson, D. N. Hayes, C. M. Perou, J. S. Parker, ABRA: improved coding indel detection via assembly-based realignment. Bioinformatics (Oxford, England) 30, 2813–2815 (2014).

50. K. Cibulskis et al., Sensitive detection of somatic point mutations in impure and heterogeneous cancer samples. Nature biotechnology 31, 213–219 (2013).

51. K. Wang, M. Li, H. Hakonarson, ANNOVAR: functional annotation of genetic variants from high-throughput sequencing data. Nucleic acids research 38, e164 (2010).

52. I. A. Adzhubei et al., A method and server for predicting damaging missense mutations. Nature methods 7, 248–249 (2010).

53. H. A. Shihab, J. Gough, D. N. Cooper, I. N. Day, T. R. Gaunt, Predicting the functional consequences of cancer-associated amino acid substitutions. Bioinformatics (Oxford, England) 29, 1504–1510 (2013).

54. C. Dong et al., Comparison and integration of deleteriousness prediction methods for nonsynonymous SNVs in whole exome sequencing studies. Hum Mol Genet 24, 2125–2137 (2015).

55. C. Shyr et al., FLAGS, frequently mutated genes in public exomes. BMC Med Genomics 7, 64 (2014).

56. M. S. Lawrence et al., Mutational heterogeneity in cancer and the search for new cancer-associated genes. Nature 499, 214–218 (2013).

57. J. Ou, L. J. Zhu, trackViewer: a Bioconductor package for interactive and integrative visualization of multi-omics data. Nature methods 16, 453–454 (2019).

58. E. Talevich, A. H. Shain, T. Botton, B. C. Bastian, CNVkit: Genome-Wide Copy Number Detection and Visualization from Targeted DNA Sequencing. PLoS Comput Biol 12, e1004873 (2016).

59. S. Nik-Zainal et al., The life history of 21 breast cancers. Cell 149, 994–1007 (2012).

60. C. H. Mermel et al., GISTIC2.0 facilitates sensitive and confident localization of the targets of focal somatic copy-number alteration in human cancers. Genome Biol 12, R41 (2011).

61. T. Rausch et al., DELLY: structural variant discovery by integrated paired-end and split-read analysis. Bioinformatics (Oxford, England) 28, i333–i339 (2012).

62. B. J. Girard et al., Cytoplasmic PELP1 and ERRgamma protect human mammary epithelial cells from Tam-induced cell death. PLoS One 10, e0121206 (2015).

63. W. A. Braden, A. K. McClendon, E. S. Knudsen, Cyclin-dependent kinase 4/6 activity is a critical determinant of pre-replication complex assembly. Oncogene 27, 7083–7093 (2008).

64. J. F. Conklin, J. Baker, J. Sage, The RB family is required for the self-renewal and survival of human embryonic stem cells. Nat Commun 3, 1244 (2012).

65. L. Micallef, P. Rodgers, eulerAPE: drawing area-proportional 3-Venn diagrams using ellipses. PLoS One 9, e101717 (2014).

66. S. Kurtenbach, J. W. Harbour, SparK: A Publication-quality NGS Visualization Tool. bioRxiv, (2019).

67. G. Yu, L. G. Wang, Q. Y. He, ChIPseeker: an R/Bioconductor package for ChIP peak annotation, comparison and visualization. Bioinformatics (Oxford, England) 31, 2382–2383 (2015).

68. J. R. Conway, A. Lex, N. Gehlenborg, UpSetR: an R package for the visualization of intersecting sets and their properties. Bioinformatics (Oxford, England) 33, 2938–2940 (2017).

69. T. Stuart et al., Comprehensive integration of single cell data. bioRxiv, 460147 (2018).

70. A. Butler, P. Hoffman, P. Smibert, E. Papalexi, R. Satija, Integrating single-cell transcriptomic data across different conditions, technologies, and species. Nature biotechnology 36, 411–420 (2018).

71. C. S. McGinnis, L. M. Murrow, Z. J. Gartner, DoubletFinder: Doublet Detection in Single-Cell RNA Sequencing Data Using Artificial Nearest Neighbors. Cell Syst 8, 329–337.e324 (2019).

72. E. Becht et al., Dimensionality reduction for visualizing single-cell data using UMAP. Nature biotechnology, (2018).

73. V. D. Blondel, J.-L. Guillaume, R. Lambiotte, E. Lefebvre, Fast unfolding of communities in large networks. Journal of Statistical Mechanics: Theory and Experiment 2008, P10008 (2008).

