## Supplementary Data for "RB1 loss triggers dependence on ESRRG in retinoblastoma"

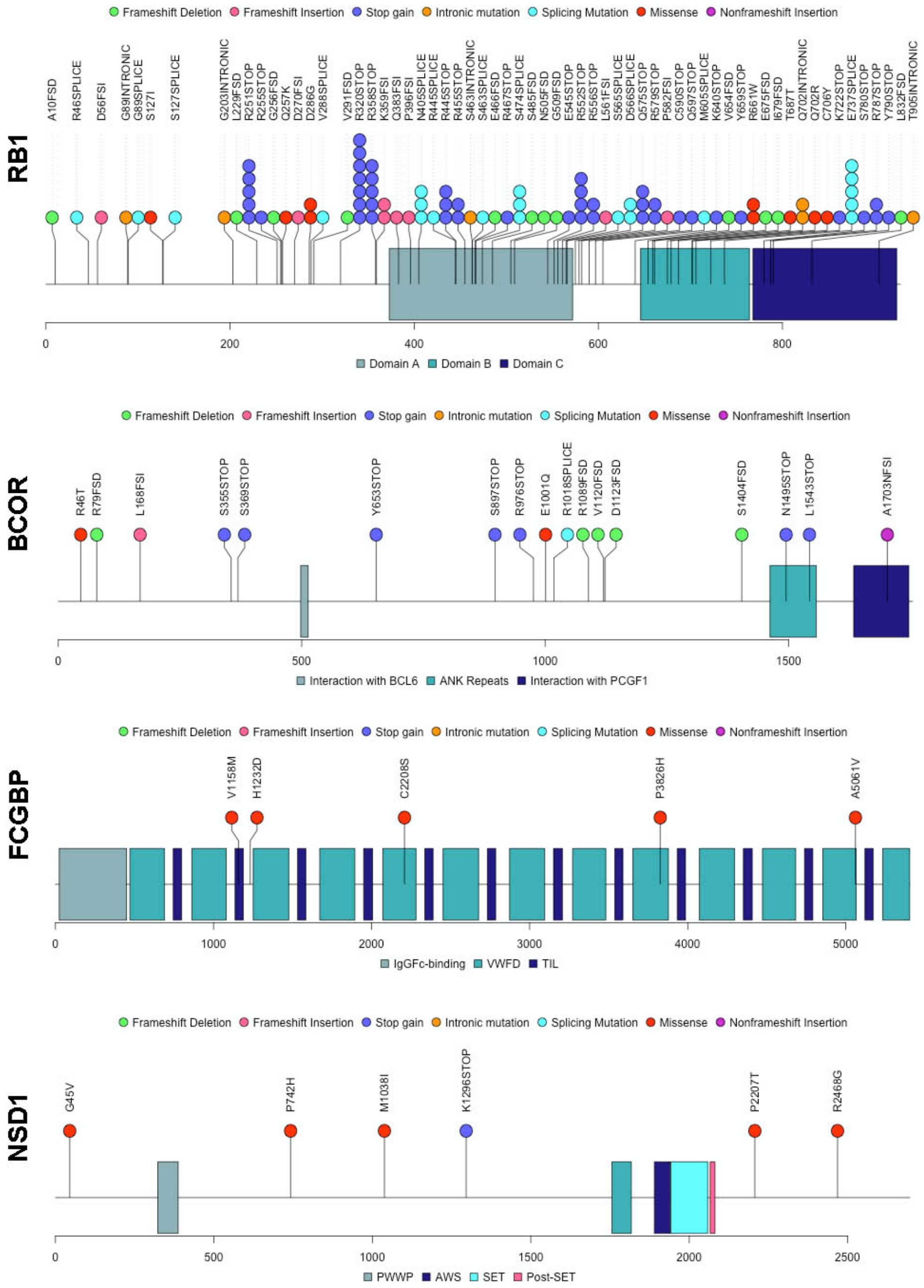


**Fig. S1. Lollipop plots of most recurrent gene mutations in 103 whole exome sequencing retinoblastoma samples.** Mutations identified in RB1, BCOR, FCGBP, and NSD1 are plotted along the protein domains and labeled by mutation type.

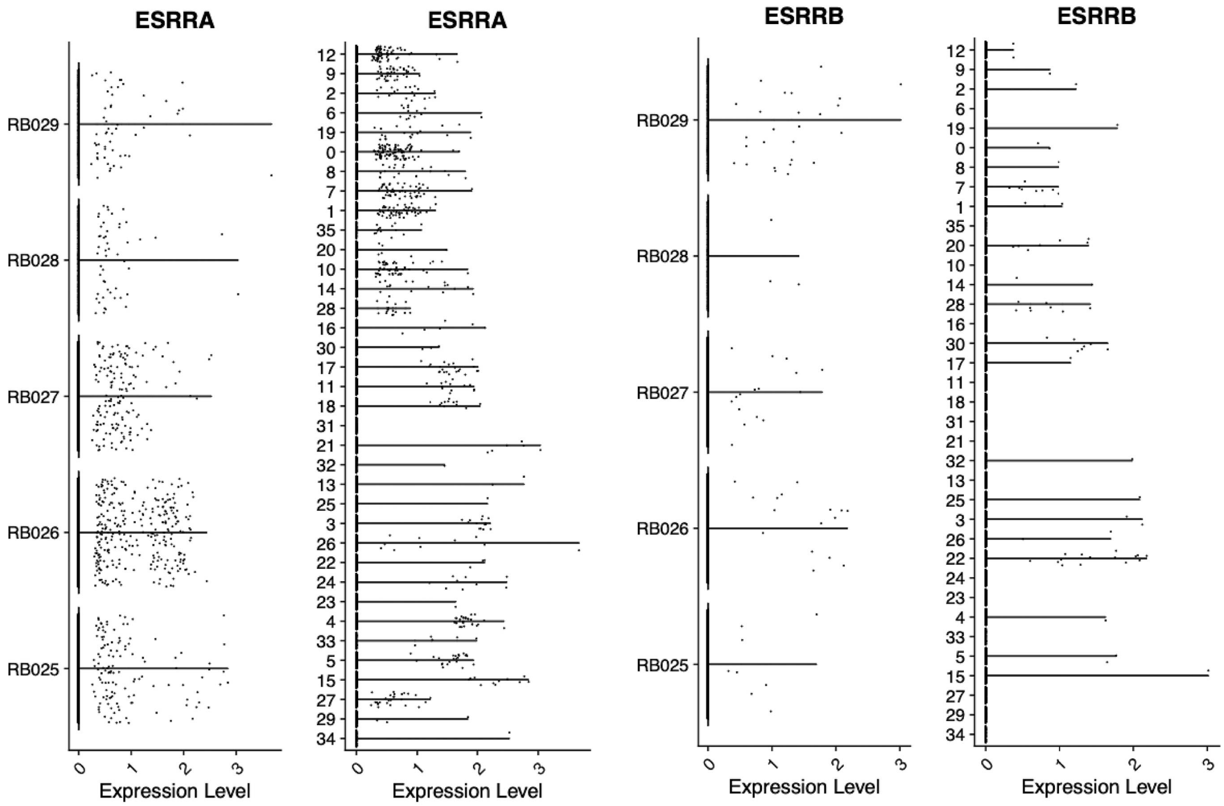


**Fig. S2. Single-cell analysis of ESRRA and ESRRB.** Violin plots of single-cell RNA sequencing (scRNA-seq) data showing expression of ESRRA (**A**) and ESRRB (**B**) by tumor sample (left) and Seurat cluster (right).

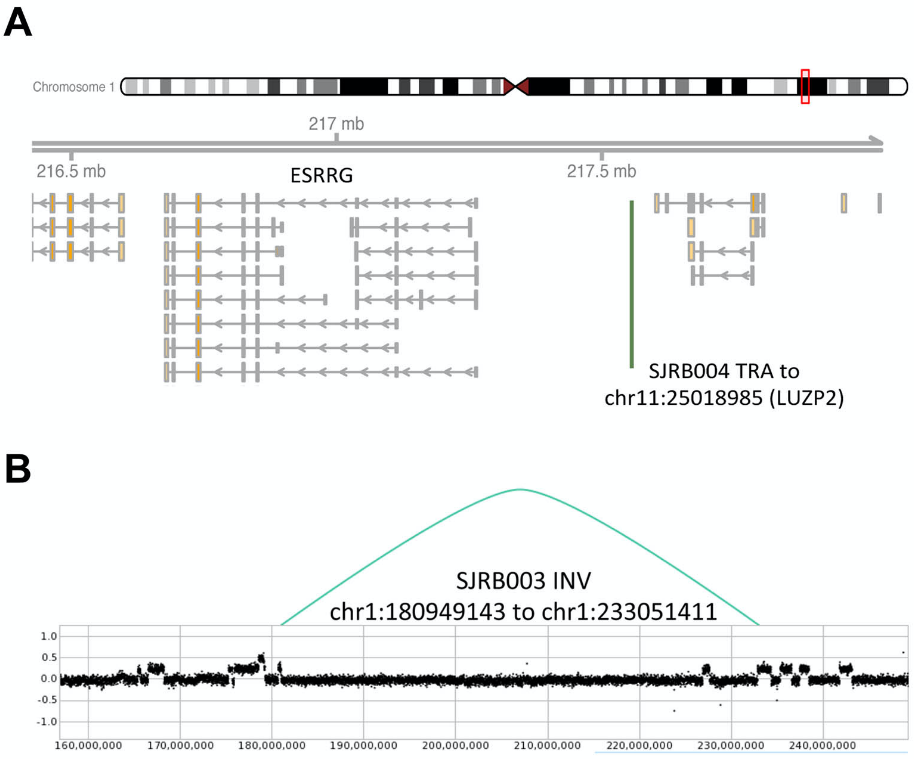


**Fig. S3. Identification of complex genomic alterations involving the ESRRG locus in whole genome sequencing data from primary retinoblastomas.** (**A**) Translocation of ESRRG (chr1:217532626) to the LUZP2 locus on chromosome 11 (chr11:25018985) in retinoblastoma sample SJRB004. (**B**) Chromosome 1q inversion (chr1:180949143 to chr1:233051411) involving the ESRRG locus in retinoblastoma sample SJRB003.

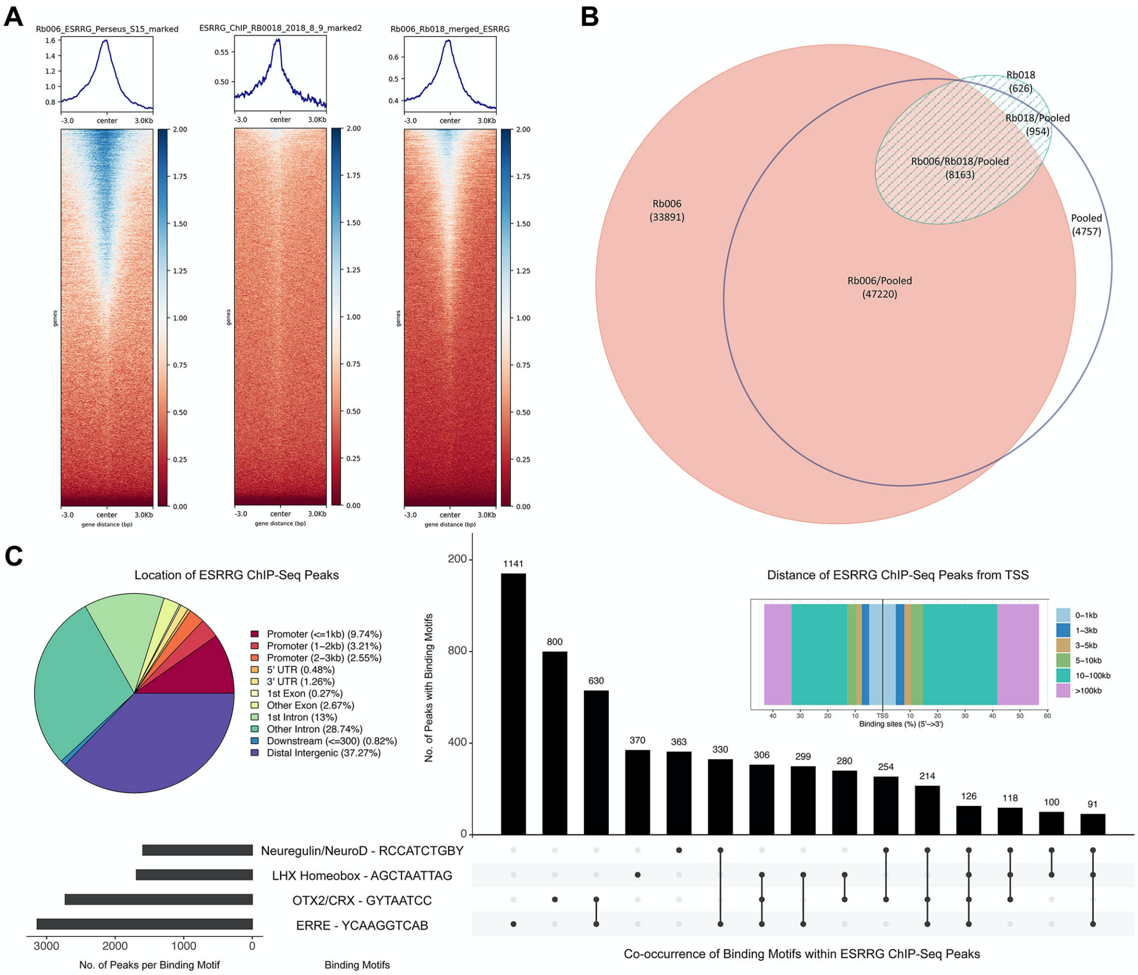


**Fig. S4. ESRRG chromatin localization in retinoblastoma cells.** (**A**) Heatmaps of called peaks from ChIP-seq pulldown of ESRRG in primary low passage RB006 and RB018 retinoblastoma cells and the two datasets pooled. Peaks ±3kb from transcription start sites (TSS) of coding genes are displayed. (**B**) Euler diagram of overlapping ESRRG ChIP-seq peaks between the 3 datasets. (**C**) Location and motif analysis of the 8163 significantly enriched ($p < 0.001$) ESRRG ChIP-seq peaks shared in common between the 3 datasets. The pie chart displays the percentage of these peaks located within various gene regions. The rectangular plot exhibits the distance of the peaks from the TSS. The bar plot shows the presence and co-occurrence of the most significantly enriched transcription factor binding motifs (FDR < 0.05) found within the ESRRG ChIP-Seq peaks.

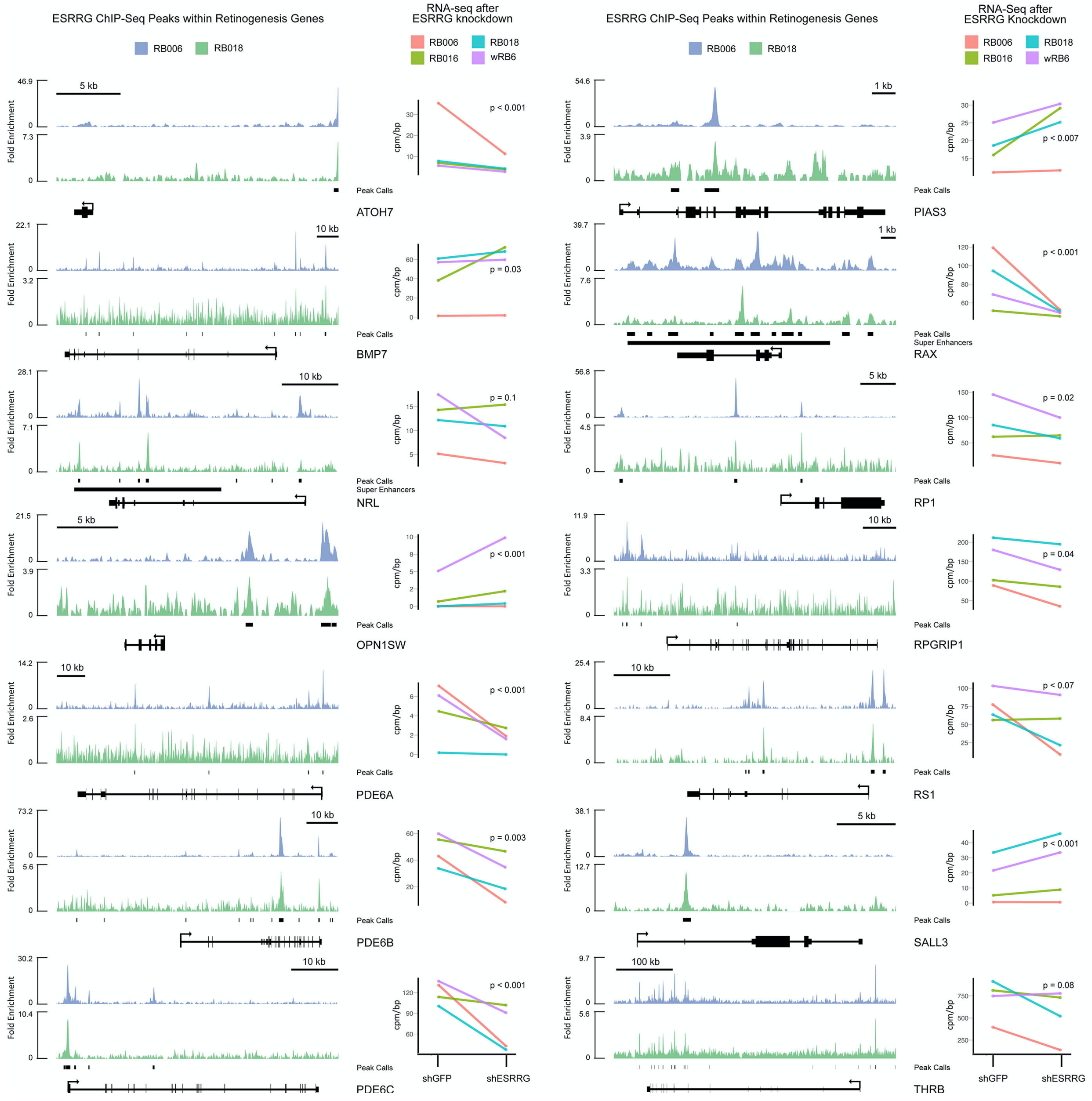


**Fig. S5. ESRRG regulates genes involved in retinogenesis.** ChIP-seq track plots from ESRRG-pulldown in RB006 (blue) and RB018 (green) retinoblastoma cells are shown for key genes involved in retinal development and differentiation. Peak calls from the pooled dataset, super enhancers, exon and intron locations, and direction of transcription are indicated below the peak plots. Corresponding RNA-seq data for each gene are shown in RB006 (red), RB016 (green), RB018 (blue), and wRB6 (purple) retinoblastoma cells engineered to express shRNA directed against ESRRG (shESRRG) or control (shGFP). P-values were calculated after batch and dispersion correction using EdgeR.

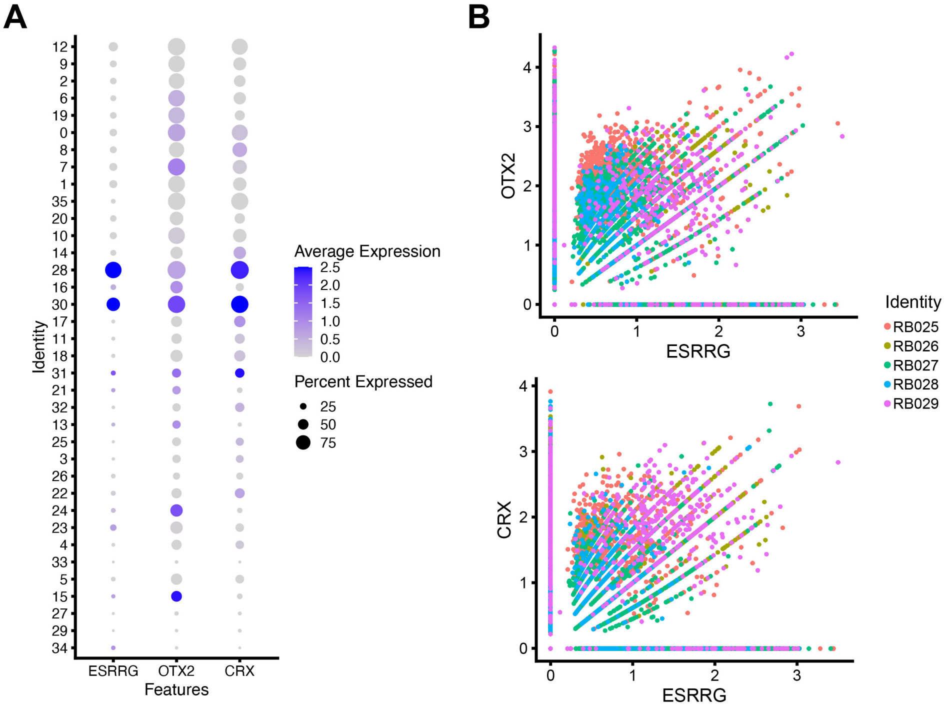


**Fig. S6. Correlation of ESRRG expression with CRX and OTX2 in single retinoblastoma cells.** (**A**) Dot plot of cell clusters described in Fig. 2C, indicating mRNA expression of *ESRRG*, *CRX* and *OTX2*. Asterisks indicate significant up-regulation, defined as log2 fold change > 0.5 and adjusted p-value < 0.05. (**B**) Scatter plots comparing expression of ESRRG to OTX2 (top) and CRX (bottom).

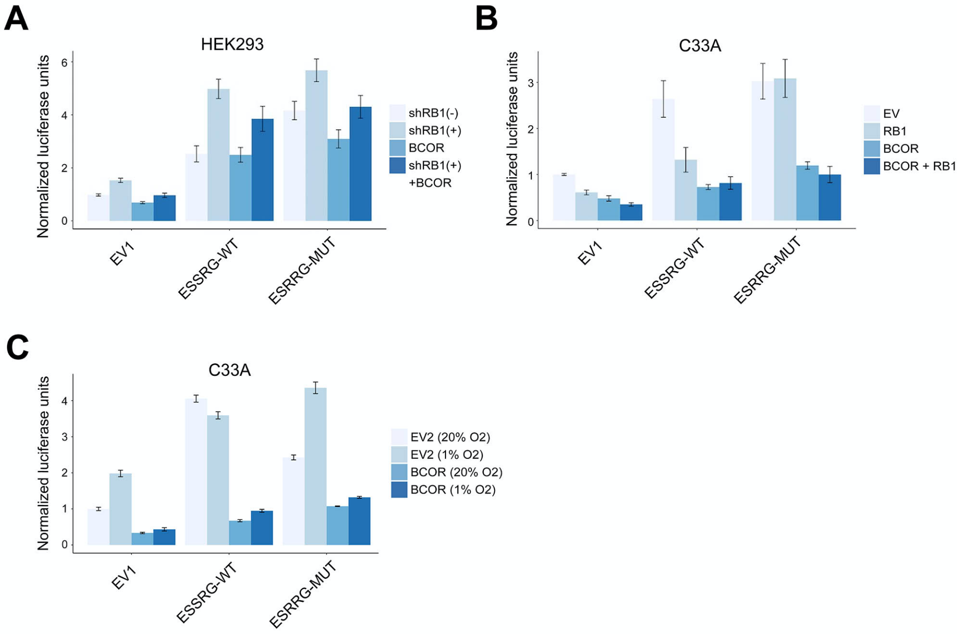


**Fig. S7. BCOR represses ERRE promoter independently of RB1.** (**A**) Normalized ERRE luciferase reporter activity with or without ectopic expression of V5-ESRRG-WT, V5-ESRRG-MUT, and BCOR, and with or without shRNA-mediated depletion of RB1 (shRB1+ and shRB1-, respectively) in HEK293 cells. (**B**) Normalized ERRE luciferase reporter activity with or without ectopic expression of V5-ESRRG-WT, V5-ESRRG-MUT, BCOR, and RB1 in C33A cells. (**C**) Normalized ERRE luciferase reporter activity with or without ectopic expression of V5-ESRRG-WT, V5-ESRRG-MUT, and BCOR in C33A cells in hypoxia (1% O2) or normoxia (20% O2). EV, EV1 and EV2, empty vector controls.

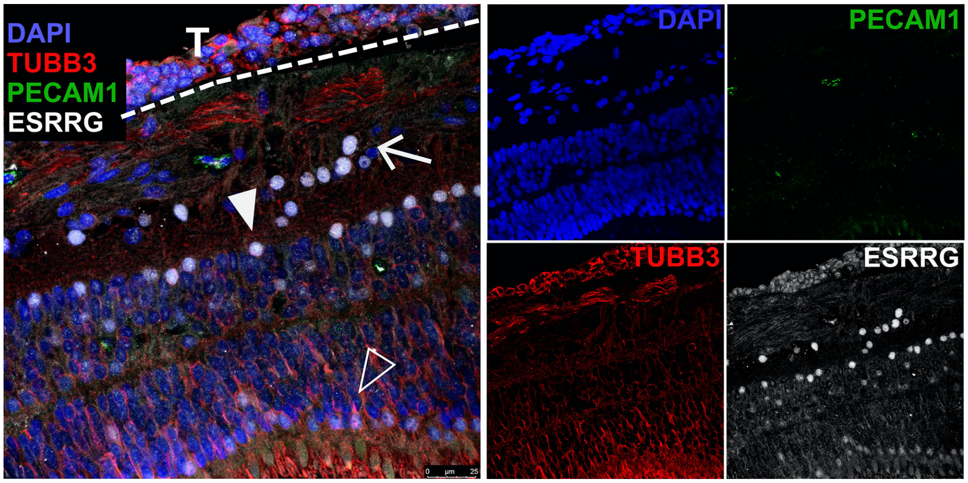


**Fig. S8. ESRRG expression in human retina.** Multiplexed fluorescence immunohistochemistry of Rb enucleation sample #29-15 in a region of unaffected retina. ESRRG (white), TUBB3 (red), DAPI (blue) and PECAM1 (green). Arrow, retinal ganglion cells; solid arrowhead, amacrine cells; hollow arrowhead, photoreceptors; T, tumor cells located on the inner retinal surface.
